# Phenotypic innovation in one tooth induced concerted developmental evolution in another

**DOI:** 10.1101/2020.04.22.043422

**Authors:** Marie Sémon, Klara Steklikova, Marion Mouginot, Manon Peltier, Philippe Veber, Laurent Guéguen, Sophie Pantalacci

## Abstract

Serial appendages are similar organs found at different places in the body, such as fore/hindlimbs or different teeth. They are bound to develop with the same pleiotropic genes, apart from identity genes. These identity genes have logically been implicated in cases where a single appendage evolved a drastically new shape while the other retained an ancestral shape, by enabling developmental changes *specifically* in one organ. Here, we showed that independent evolution involved developmental changes happening *in both* organs, in two well characterized model systems.

Mouse upper molars evolved a new dental plan with two more cusps on the lingual side, while the lower molar kept a much more ancestral morphology, as did the molars of hamster, our control species. We obtained quantitative timelines of cusp formation and corresponding transcriptomic timeseries in the 4 molars. We found that a molecular and morphogenetic identity of lower and upper molars predated the mouse and hamster divergence and likely facilitated the independent evolution of molar’s lingual side in the mouse lineage. We found 3 morphogenetic changes which could combine to cause the supplementary cusps in the upper molar and a candidate gene, *Bmper*. Unexpectedly given its milder morphological divergence, we observed extensive changes in mouse lower molar development. Its transcriptomic profiles diverged as much as, and co-evolved extensively with, those of the upper molar. Consistent with the transcriptomic quantifications, two out of the three morphogenetic changes also impacted lower molar development.

Moving to limbs, we show the drastic evolution of the bat wing also involved gene expression co-evolution and a combination of specific and pleiotropic changes. Independent morphological innovation in one organ therefore involves concerted developmental evolution of the other organ. This is facilitated by evolutionary flexibility of its development, a phenomenon known as Developmental System Drift.

**AUTHOR SUMMARY:** Serial organs, such as the different wings of an insect or the different limbs or teeth of a vertebrate, can develop into drastically different shapes due to the position-specific expression of so-called “identity” genes. Often during evolution, one organ evolves a new shape while another retains a conserved shape. It was thought that identity genes were responsible for these cases of independent evolution, by enabling developmental changes specifically in one organ. Here, we showed that developmental changes evolved *in both* organs to enable the independent evolution of the upper molar in mice and the wing in bats. In the organ with the new shape, several developmental changes combine. In the organ with the conserved shape, part of these developmental changes are seen as well. This modifies the development but is not sufficient to drastically change the phenotype, a phenomenon known as “Developmental System Drift”, DSD. Thus, the independent evolution of one organ relies on concerted molecular changes, which will contribute to adaptation in one organ and be no more than DSD in another organ. This concerted evolution could apply more generally to very different body parts and explain previous observations on gene expression evolution.

## INTRODUCTION

Serial appendages are repetitions of similar appendages in the body, such as different legs or wings in arthropods, vertebrate fore- and hindlimbs, or different teeth. According to their position and function, they can have very similar or very different shapes. Although a certain degree of individuation is often present deeply in evolution, its magnitude can evolve, with new shapes appearing specifically in one appendage. For example, fore- and hindlimb have had different shapes since early tetrapod evolution (Minelli, 2003; Siomava et al., 2020). In bats, the forelimb evolved into a wing, while the hindlimb conserved a more ancestral morphology (Cooper et al., 2012; Sadier et al., 2021). Similarly, in insects, one of the two pairs of wings was modified to form elytra and haltere in coleoptera and diptera, respectively (Tomoyasu, 2017). How selection could act to drastically change the shape of one serial appendage independently of another is an intriguing question.

The reason for that is the pleiotropy constraint. During development, serial organs develop with the same pleiotropic genes, except a handful of key transcriptional regulators, whose expression is appendage-specific. These “selector genes” or “identity genes”, including the famous homeotic genes, are necessary to form the right appendage at the right place (Mann & Carroll, 2002; Tomoyasu, 2017; Weatherbee & Carroll, 1999). For example, *Ubx* is necessary to form a haltere instead of a wing by regulating hundreds of pleiotropic target genes specifically in the haltere (Hersh et al., 2007; Pavlopoulos & Akam, 2011)). The enhancers of these target genes are surprisingly pleiotropic as they are shared with the wing (McKay & Lieb, 2013). Intuitively, mutations in identity genes, or in the regulatory regions they target, might be easily selected because they will inherently have appendage-specific effects (Carroll, 2008; Morgalev et al., 2023). In contrast, mutations in other parts of pleiotropic genes might more often be counterselected, because they may have an effect on development of both appendages, which may be advantageous in one appendage but deleterious in the other. Therefore, there is an expectation that identity genes will play a central role in the independent evolution of serial appendages.

Many examples of independent appendage evolution have been studied, which confirm the role of identity genes, but not only.

In the simplest cases, just a subpart of the appendage has changed, such as hairless parts or specialized hair structures (eg. sexcombs) in fly legs. As expected, these new traits were associated with new expression patterns of homeotic genes specific of one leg (eg. *Ubx*; *Scr*) or their targets (eg. *Dsx*) and with the evolution of their cis-regulatory regions (G. K. Davis et al., 2007; Eksi et al., 2018; Stern, 1998; Tanaka et al., 2011) . This nicely explains why these fly leg structures evolved at a specific position (or in one sex), but the developmental and genetic changes through which selection has shaped these structures is still under study (Atallah et al., 2014).

More complex cases concern the whole appendage, such as halteres and elytra in insects (Tomoyasu, 2017), jump-adapted legs in insects (Mahfooz et al., 2007; Refki et al., 2014) and rodents (Saxena et al., 2022), wings in bats (Cooper et al., 2012; Sadier et al., 2021), or patterns of eyespots in butterflies fore- and hindwings (Matsuoka & Monteiro, 2021, 2022). Here again studies have pointed to a role for homeotic and identity genes in general (Booker et al., 2016; Matsuoka & Monteiro, 2022; Refki et al., 2014; Saxena et al., 2022). Moreover, in species where wings are well differentiated (e.g. wing/haltere of flies, or to a lesser extent, anterior/posterior wings of bees), the overall dose of hox genes appears more different between the two appendages than in species with less differentiated appendages (e.g. anterior/posterior wing of dragonflies), suggesting that the evolution of a differential hox dose is an important determinant of appendage differentiation (Paul et al., 2021). Other developmental genes have been implicated as well, because they evolved a new expression pattern in the modified appendage that is consistent with its phenotype. (Saxena et al., 2022; Z. Wang et al., 2014). More intriguingly, transcription factors which are expressed in both appendages have appendage-specific functions revealed by knock-out experiments or tests in heterologous species (Cretekos et al., 2008; Matsuoka & Monteiro, 2022; Ravisankar et al., 2016; Tomoyasu et al., 2009). It is assumed that this appendage-specific function is provided by identity genes, either directly (through unknown differential cis-regulation in the two appendages) or indirectly by providing context-dependency, but this remains generally untested. Together this indicates that the independent evolution of serial organs does cope with pleiotropic genes, but does not fully explain how.

Here we chose a new model and a different approach to address this question, focusing on developmental dynamics. We studied the independent evolution of a drastically new shape in the mouse upper molar, which is nicely described in the fossil record. Since it occurred relatively recently, we could compare closely related species. Another strong advantage of this model is that the development of mouse molars is very well understood mechanistically, from years of developmental genetics and morphogenesis modeling. Finally, we devised specific ways to quantify and decipher the evolution of development based on the comparison of transcriptome time series.

Between 18-12 million years ago, the upper molars of mouse and rat ancestors gradually acquired a new cusp row on the lingual side of the molar, and reduced cusps size on the buccal side (Figure 1). This new dental plan and accompanying changes in mastication movements are adaptive and associated with the success of murine rodent radiation (Lazzari et al., 2008; Tiphaine et al., 2013). Changes in the lower molar were limited to the connections between cusps, keeping cusp number and size constant (Figure 1). Because the shape of upper and lower molars were different although less individuated in the basal “cricetine” rodents, from which murine rodents emerged (Figure 1) this is not a case of *de novo* individuation. In fact, lower and upper molars already had different morphologies in the first mammals (B. M. Davis, 2011; Hillson, 2005). Hamsters are today’s good representative of the basal “cricetine” rodents. In the golden hamster lineage, both molars kept the ancestral cusp number and organization. We can make the reasonable assumption that the hamster presents ancestral developmental features, and in this study we use this species as a phylogenetic control.

**Fig 1.**
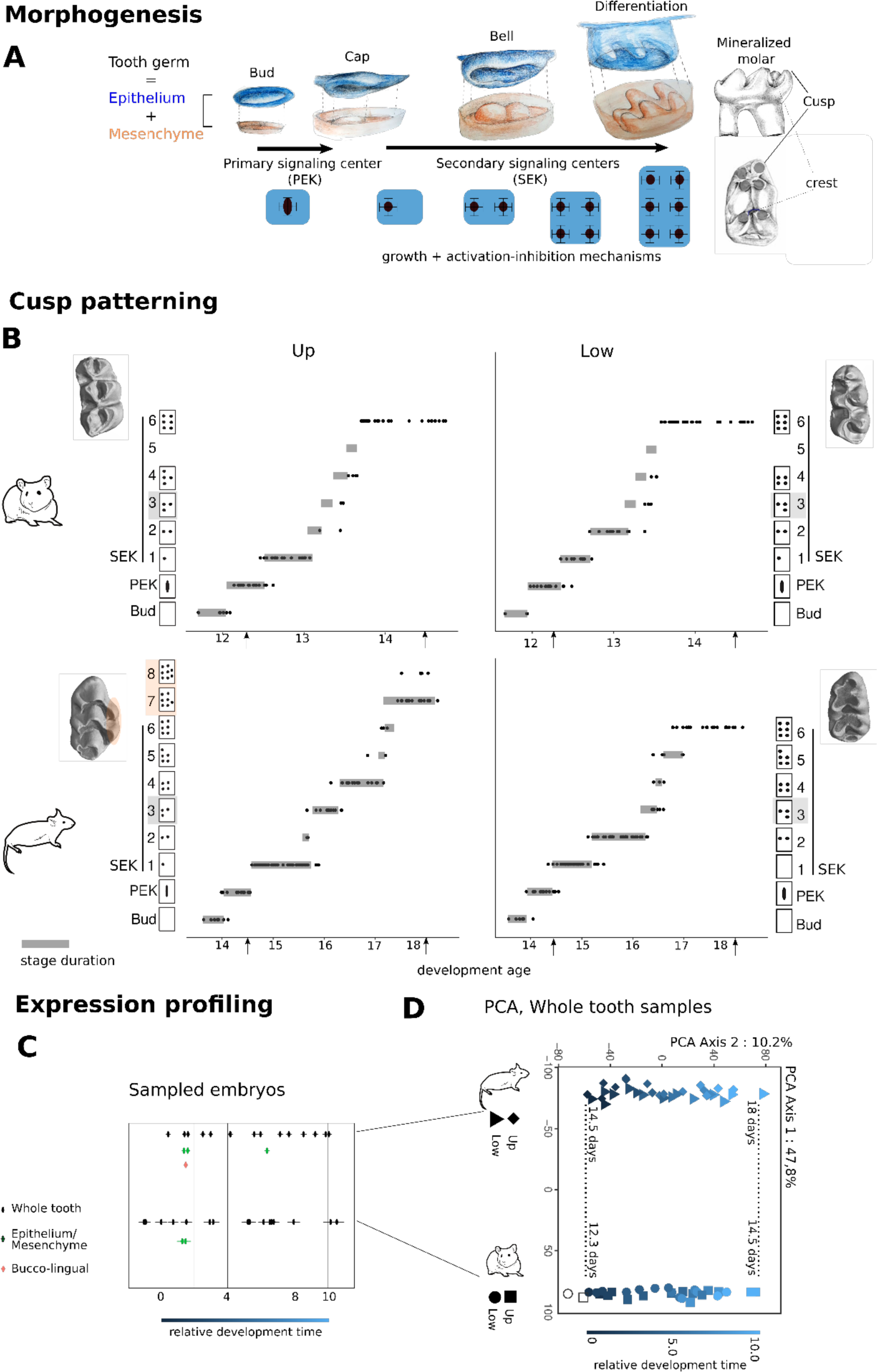
Comparing molar development in different tooth types and species. A. Drawings of the epithelium and mesenchyme compartments at four stages of molar development: at “bud” stage, an epithelial signalling centre called PEK (for Primary Enamel Knot) triggers the formation of a “cap” defining the future crown. Between “cap” and “bell” stage, SEKs (Secondary Enamel Knots) are patterned sequentially in the epithelium, and drive cusp formation. By the end of morphogenesis, the mesenchyme has the shape of the future tooth and the epithelium is a kind of dental impression. B. Dynamic and pattern of PEK/SEK addition in mouse and hamster lower and upper molars. Each panel is a series of developing molars hybridised against Fgf4 to reveal signalling centres (back dots). x axis: developmental age, y axis: morphological stage, with diagrams of signalling centres arrangements and 3D scans for final morphologies. Time series were modelled using Markov processes as a series of stages with specific durations (grey bars). Arrowheads show homology established on morphological criteria at early cap stages and late bell stage. C. Embryo sampling used for expression profiling, in whole tooth germs, tooth tissues (mesenchyme/epithelium) and tooth halves (bucco/lingual). Each embryo provided upper and lower samples. Relative developmental time established from embryonic weight and boundaries for stage homology as in D. D. Principal component (PC) analysis of 64 whole molar bulk transcriptomes based on 14532 1:1 orthologues. Each symbol is an individual transcriptome with a colour gradient for relative development time. Dotted lines: Stage homology established by morphology is confirmed and used as boundaries for relative developmental time.

Molars develop from the physical and molecular interaction between an epithelium and a mesenchyme (Jernvall & Thesleff, 2012), Figure 1A). The epithelium grows and folds to form the crown and its cusps under the influence of two types of signalling centres, PEK and SEK (Primary and Secondary Enamel Knots respectively) (Jernvall & Thesleff, 2012). First, the PEK determines the field of the molar crown. As this field grows, the SEKs are patterned sequentially and determine the cusps, starting with a buccal cusp (Cho et al., 2007; Pantalacci et al., 2017). This spatio-temporal sequence depends on activation-inhibition loops involving both epithelium and mesenchyme in a Turing-like mechanism (Salazar-Ciudad, 2012). Tooth morphogenesis models and *in vivo* experiments have shown that changes in the pathways controlling these loops can modify the number of cusps and recapitulate evolutionary changes (Harjunmaa et al., 2012, 2014; Morita et al., 2020; Salazar-Ciudad & Jernvall, 2010). More minimal modeling of Turing-like mechanisms in teeth has shown how the interaction between activation-inhibition loops and growth of the field dynamically determines the output pattern (Morita et al., 2022; Sadier et al., 2019) . Applied to the case of supplementary lingual cusps, this theoretical framework predicts that the new phenotype could be achieved by a change in the activation-inhibition loops (e.g. allowing cusps forming closer from each other), a change in bucco-lingual (B/L) growth (allowing more cusps to be fitted in a bigger field), or a combination of both.

Almost all what is known was established on the lower first molar of the mouse and few studies have compared lower and upper molar development. The homeotic-code present in the jaws at the early stages of development is still present when molars are initiated (*Dlx1/2* and *Pou3f3* for upper jaw, *Dlx1/2/5/6* and *Nkx2.3* for lower jaw (Cobourne & Sharpe, 2003; Hirschberger et al., 2021; Jeong et al., 2008). We have previously demonstrated that later during morphogenesis, the expression of *Dlx5/6* genes is no longer specific but remains biased (Pantalacci et al., 2017). Only three genes remain specific: in the upper molar, *Pou3f3* and its non-coding regulator and in the lower molar, *Nkx2.3*. On top of that, many genes are consistently biased throughout morphogenesis (Pantalacci et al., 2017). The genetic architecture of lower and upper molars differs in mouse, since a few mouse mutants have molar-specific phenotypes (including Dlx1/2 and Pitx1; reviewed in (Hallikas et al., 2021; Kwon et al., 2017)), and molar specific loci are evidenced in the two available quantitative genetic studies (Navarro & Murat Maga, 2018; Shimizu et al., 2004).

On top of working with this well characterized system, we took a comparative analysis of RNA-seq timeseries, which proved successful to study development and its evolution. RNA-seq temporal profiles describe the dynamic changes which orchestrate the development of a complex structure, such as the many transcriptional and cell proportions changes (Pantalacci & Sémon, 2015). In our published RNAseq timeseries comparing the development of the mouse lower and upper first molar, we found that differences in their transcriptomes were reflecting differences in their morphogenesis, such as different proportions in the respective tissues of the tooth or different rates of cusp formation in the tooth (Pantalacci et al., 2017). This showed that transcriptome timeseries contain valuable information to compare the morphogenesis of two organs, even though this information is not immediately noticeable. Comparing the transcriptomes of different species provides a useful quantification of developmental similarity - once possible biases in estimating expression levels have been controlled for (Cardoso-Moreira et al., 2019; Pantalacci & Sémon, 2015). This has been successfully applied to answer questions on the periods of maximal conservation of embryogenesis (so-called hourglass pattern, eg (Kalinka et al., 2010; Levin et al., 2016)) or on the homology of organs (Fisher et al., 2020; Tschopp et al., 2014; Z. Wang et al., 2011).

Here, to understand how lower and upper molars evolved independently from one another, we compared the dynamics of their developmental systems in mouse and hamster with a twofold strategy, analyzing 1) RNA-Seq time series and 2) the dynamics of cusp formation, obtained by tracking a cusp marker in hundreds of molar samples. This revealed an ancestral molecular identity for each molar type, associated with morphogeneticspecificities. Consistent with the very peculiar mouse upper molar morphology, the two mouse molars have more different temporal profiles than the two hamster’s. We found three morphogenetic changes which could combine to cause the supplementary cusps in the upper molar, one of them building on the ancestral specificity of upper molars. The biggest surprise came from the lower molar, which was first thought of as an additional control. As many gene expression temporal profiles diverged in the lower molar as in the upper molar, and a great part of them are co-evolving in the two molars. This is associated with changes in B/L polarity and activation-inhibition mechanisms of cusp formation seen in both molars. Based on these results, we propose that several mutations combined to reach the new upper molar dental plan, some with specific developmental effects, and others with effects in both molars. As a consequence, drift in lower molar development went along with adaptation in upper molar development. We generalised our findings by re-analysing transcriptome and literature data on bat foot and wing evolution. We propose that mutations producing shared gene expression changes have a major contribution to appendage-specific adaptation.

## RESULTS

Mouse and hamster molars develop at different paces. We predicted developmental age from embryonic weight in each species and aligned temporal series between species with homologous start and end points of first molar morphogenesis (Figures 1 and S1). We then devised a twofold strategy to compare the dynamics of cusp formation along with the dynamics of gene expression. First, we established the sequence of PEK and SEKs formation and modelled the relative stage durations with continuous Markov processes (Figure 1B). Second, we obtained RNA-seq data from whole first molar germ at high time resolution (Figure 1C) to model temporal profiles and added samples for dissociated epithelium and mesenchyme as well as buccal and lingual half germs for specific purposes (Figure 1C).

### Conserved transcriptomic and morphogenetic features point to an ancestral identity of lower and upper molars

Before focusing on the independent evolution of the first upper molar in mouse, we first looked for molecular and developmental features which discriminate between lower and upper molar development in the two species. This would form a molecular and developmental identity of the molars common to both species, and likely present in their common ancestor. It could have served as a basis for the independent evolution of the upper molar in the mouse lineage.

#### Identity genes and many other key genes for tooth morphogenesis showed a conserved expression bias that distinguish lower and upper molars in both species

Principal components in a PCA analysis separate samples according to the main axes of variation in the data. In our dataset, the main effect was the species, followed by development time (Figure 1C). Upper and lower molar samples were only separated from each other on the sixth component, which carried 3% of the total variance. Hence only a minor part of the variation distinguishes upper and lower samples while being common to both species.

To detect genes carrying this conserved variation between upper and lower molars, we modelled their temporal profiles with polynomials (Figure 2A). In each species, we fitted two models: one with two distinct curves, one per tooth, and another with a single curve, common for both teeth. By comparing the fit of these two models, we detected differentially expressed genes between the two teeth (Table S2). In each species, we found a similar proportion of differentially expressed genes (about 13%). There are 712 genes in common between the two species (only a third of the genes in each species), including 550 genes with a consistent upper/lower bias.

**Fig 2.**
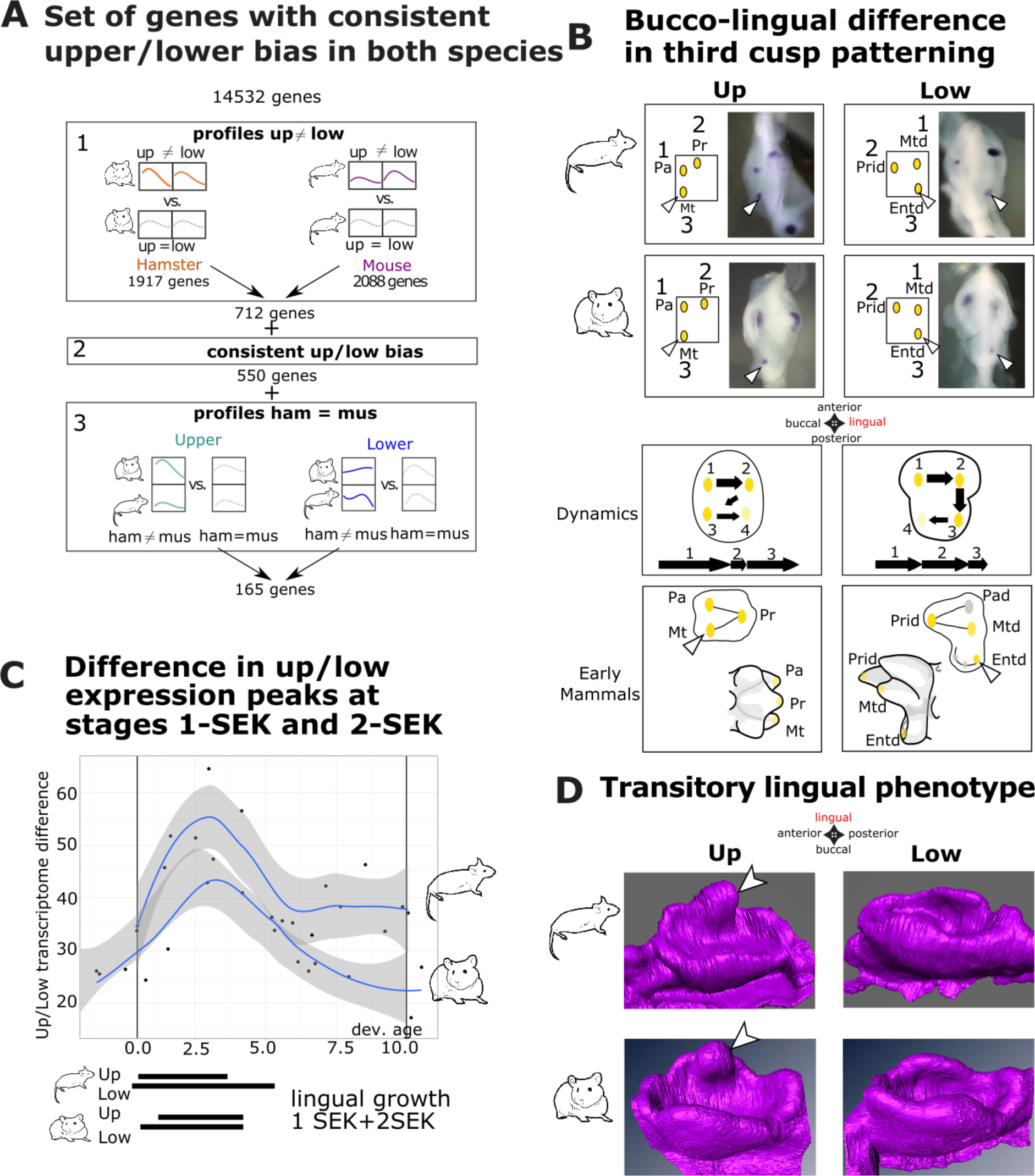
Conserved differences of upper and lower molar development. A. Pipeline to search for genes that distinguish lower and upper molars in the two species. 1/ Expression profiles were modelled separately in hamster (left) and mouse (right) to detect genes with distinct profiles in upper and lower molars in each species. Colored curves are models allowing for distinct profiles in upper and lower molars. Grey curves allow a single profile, common for both teeth. The resulting number of differentially expressed genes is shown. 2/ Filtering for a consistent upper/lower bias in both species. 3/ Filtering for conserved expression profiles in both species by modelling separately upper (left) and lower (molars). Colored curves: with distinct profiles in mouse and hamster. grey curve: single profile. B. Bucco-lingual difference in cusp patterning. The 3-SEK pattern involves a lingual cusp (Entd) in the lower molar and a buccal cusp (Mt) in the upper molar (arrowheads). Images show the epithelium at the 3-SEK stage, hybridized with a *Fgf4* probe. The dynamics of cusp formation is also specific to upper and lower molars. Both are reminiscent of the bucco-lingual pattern in early mammals’ tribosphenic molars (drawn from top and side views). Protoconid-Prid, Metaconid-Mtd, Entoconid - Entd, Paraconid-Pad (no homologous cusp in rodent’s molars), Paracone-Pa, Protocone-Pr, Metacone-Mt. C. The development of upper and lower molars differ mostly at early/mid-morphogenesis. The axis produced by BCA (Between Component multivariate Analysis) captures a variation between upper and lower molar that is common to both species. Each dot represents the variation measured for one individual. Peaks correspond to stages 1-SEK and 2-SEKs in the 4 molars (black bars taken from stage duration in Figure 1). D. Transitory phenotype in upper molars. At the end of the cap stage, upper molars of both species show an epithelial bulge (arrowhead) never seen in the lower molar.

The set of 550 genes with significant and consistent upper/lower bias in both species is highly relevant: it contains the expected jaw-identity genes known for mouse (*Nkx2.3*, *Pou3f3*, *Dlx1*, but also *Dlx5* and *6* which are no longer lower-jaw specific at this stage, but show a lower molar bias as already described in mouse (Pantalacci et al., 2017)), a fifth of the genes whose mutant shows a phenotype in the lower molar (21 out of 87 “keystone genes” from (Hallikas et al., 2021)) and key transcriptional regulators of molar morphogenesis (*Barx1*, *Msx1*, *Pitx1* Figure S2). Overall, these genes are strongly enriched for transcriptional regulators. They are involved in epithelial and mesenchymal development, cell adhesion, proliferation and differentiation and signalling, especially WNT, BMP and NOTCH (enrichment for Gene Ontology terms, Figure S3). Among the 550 genes with a consistent upper/lower bias, only 165 display evolutionary conserved temporal dynamics in both teeth (stars Figure S2, Table S2).

#### The earliest phase of cusp patterning gathers many features that distinguish lower and upper molars in both species

We next looked for criteria that would distinguish cusp patterning dynamics in lower and upper molar in both species, and may have been conserved from the common ancestor of mouse and hamster.

First, we noticed that in both species, SEK formation is initiated (transition to 1-SEK stage and 2-SEK stage) and completed (transition to 6-SEK stage) later in the upper molar as compared to the lower molar of the same embryo. This delay was also obvious in the transcriptome: the upper molar transcriptome looks “younger” than the lower molar transcriptome of the same embryo (Figure S4).

Second, we examined the details of the sequence of cusp acquisition. The first and second SEK are homologous in all teeth, the first SEK being buccal, and the second being its lingual neighbor. This bucco-lingual sequence is thus similar in lower and upper molars in both species, as previously shown in mouse (Cho et al., 2007; Pantalacci et al., 2017). The 3-SEK stage however distinguishes lower and upper molars in both species, with a buccal third SEK in upper molars, and a lingual third third SEK in lower molars (Figures 1B, 2B). It is striking that the three cups patterned at this 3-SEK stage recapitulate the bucco-lingual arrangement of their homologous cusps in the molars of early mammalian ancestors (Figures 2B, S5). These so-called “tribosphenic molars” were markedly asymmetric along the B/L axis (B. M. Davis, 2011; Hillson, 2005). On top of this geometric pattern, we found two other conserved distinctive features: the dynamics of the three first SEK stages (Figure 2B and figure S5), and a peculiar lingual epithelial bulge specific to the upper molar (Figure 2D). Both occur at this same early period of development. In fact this is the period where lower and upper molar transcriptomes differ most from one another, as seen on a multivariate analysis (BCA) performed on all samples to capture a lower/upper molar variation common to both species (Figure 2C).

Together, we found a clear conserved transcriptomic identity of each molar in the form of a conserved expression bias for identity genes and many key regulators of tooth development, and a conserved transcriptomic signature. A clear conserved morphogenetic identity was obvious in the earliest phase of cusp formation, with different dynamics of cusp formation along the bucco-lingual axis, that recapitulated the bucco-lingual specificities of early mammals’ molars. We next asked how the mouse upper molar evolved its new morphology.

### Increased upper-lower molar dissimilarity of mouse transcriptomes

Some transcription factors which made the ancestral identity showed increased levels and/or biases in mouse molars

Because the dose of identity transcription factors was shown to correlate with the degree of appendage differentiation, we first first asked whether the dose of transcription factors forming the conserved transcriptomic identity has changed in mouse (Figure 3A). We expected to observe an increased bias in mouse, especially in the upper molar.

**Fig 3.**
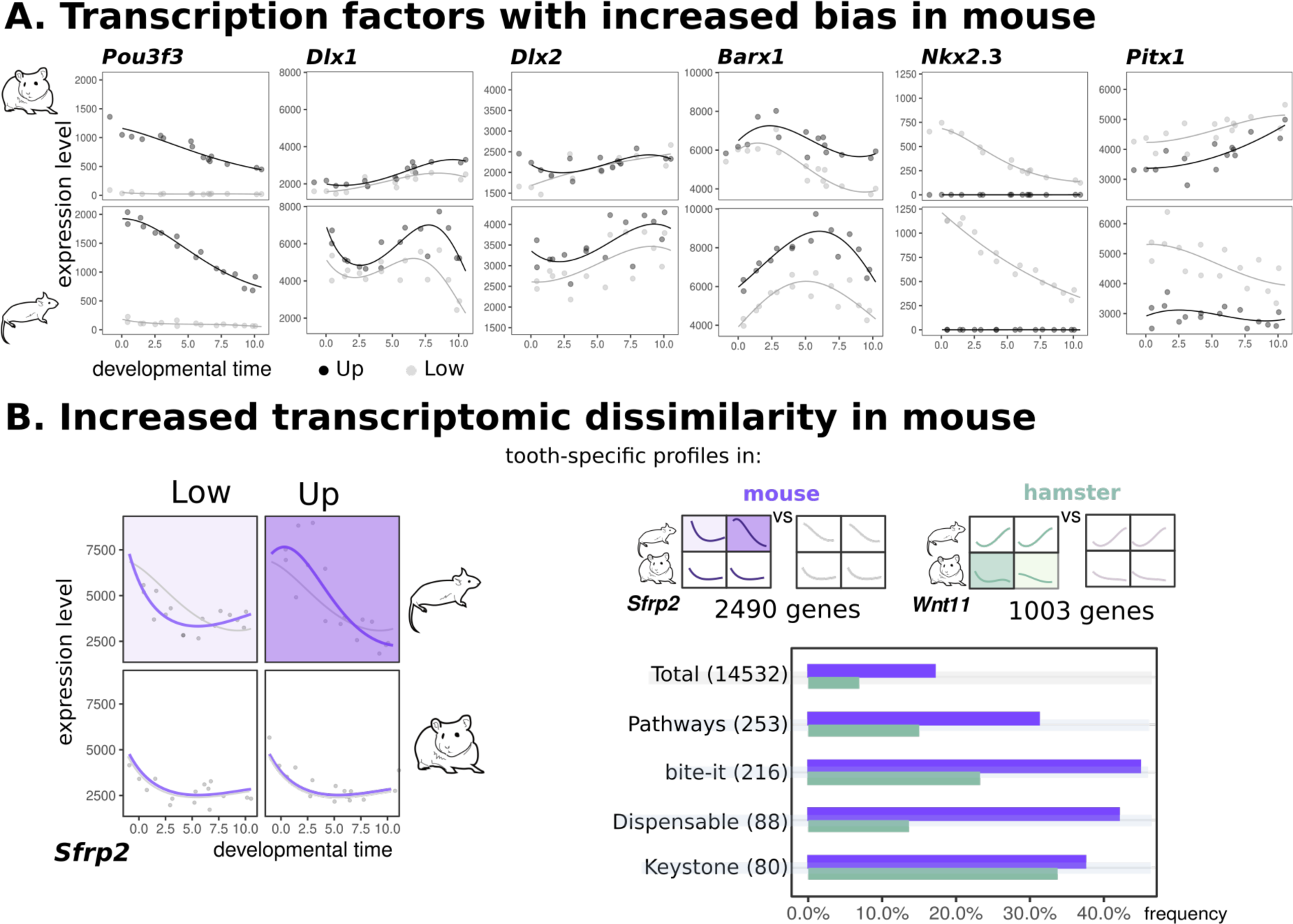
Increased upper-lower molar dissimilarity of mouse transcriptomes. A. Temporal profiles for 6 key transcription factors, distinct curves were fit for upper (black) and lower (grey) molars. B Left: Model to detect genes which differ in their upper-lower expression profiles in one species. Example of *Sfrp2* gene expression levels (grey dots), significantly better modelled by two curves in mouse and one in hamster (purple curves), compared with one curve per species (grey curves). Right: Number of genes detected by this tooth-specific model in mouse (purple) and in hamster (green, LRT with adjusted p < 0.05). Barplots show their frequencies for the “total” gene set, genes from developmental “pathways”, genes from “bite-it” database, and genes with a mild or strong phenotype in tooth mutants (“Dispensable” or “Keystone”). Size of each gene set into brackets.

*Pou3f3* is the only TF specific of the upper molar in both species. Its expression showed a twofold increase in mouse upper molar. *Dlx1/2* genes are expressed in both molars, but are essential only for upper molar formation in mouse (Qiu et al., 1997). Their expression levels were more than twice increased in mouse molars and the ratio, slightly in favor of the upper molar in hamster, is increased in mouse. *Barx1* is a key molar-specific TF whose levels have been correlated with cusp number in mammalian molars (Miletich et al., 2011). The bias in favor of the upper molar was markedly increased in mouse. This effect is selective since *Msx1*, another TF which cooperates with *Barx1 (Miletich et al., 2011)*, showed similar bias in the two species (Figure S2).

Surprisingly, such changes were not restricted to the upper molar. The expression of *Nkx2.3,* the specific TF of the lower molar, showed an almost twofold increase in mouse. For *Pitx1*, a shared TF whose mutation impairs more specifically lower molar development (Mitsiadis & Drouin, 2008), the ancestral bias in favor of the lower molar was increased. *Dlx1* expression levels were also twice increased in the lower molar. This is not true for the *Dlx5-6* genes, which were more specifically associated with lower jaw identity (Depew et al., 2005), but are expressed at this stage in the two molars (Pantalacci et al., 2017) (Figure S2). Thus, the molecular identity of each molar was reinforced in mouse, partly in line with an ancestral bias.

#### The dissimilarity of upper/lower molar transcriptomes is increased in mouse

Next, we asked whether the transcriptomes would capture a general increase of differentiation of molar development in mouse as compared to hamster. As exemplified above, a number of genes have a marginally or small significant bias in hamster, which is increased in mouse. This is in agreement with the multivariate analysis presented in Figure 2C. Along an axis that captures a lower/upper molar variation common to both species, the variation is higher for mouse than for hamster. Thus, an ancestral dissimilarity is exaggerated in mouse.

We detected genes whose expression profiles differ between upper and lower molars in mouse but not in hamster, by a dedicated model based on the 4 molars altogether (Figure 3B). As a control, we built a reciprocal model to detect genes with tooth specific profiles in the hamster. We found 2.5 times more genes with tooth-specific profiles in mouse as compared to hamster. Even after removing the effect of baseline expression levels, which may be impacted by differences in cell composition (Pantalacci et al., 2017), we still observed 1.6 times more tooth-specific profiles in mouse. The function of the genes with tooth specific profiles differs markedly between mouse and hamster (Figure S6). In mouse, genes are linked to cell adhesion and migration, as well as cell cycle and mitosis. This is consistent with our previous findings suggesting that morphogenetic movements are enhanced in mouse upper molar during the period of lingual growth (Pantalacci et al., 2017).

Thus, we quantify an increased dissimilarity of temporal profiles in mouse, consistent with the increased morphological dissimilarity of the adult teeth. This involved the reinforcement of an ancestral molecular identity as well as newly evolved gene expression differences.

### Morphogenetic changes associated with mouse upper molar evolution

Next, we used the transcriptome as a starting point to investigate several possible developmental mechanisms how the mouse upper molar forms additional lingual cusps. We logically focused on specificities of mouse upper molar development as compared to any other teeth.

#### The proportion of mesenchyme is increased in the mouse upper molar

We previously showed that the upper molar germ of the mouse contains more mesenchyme relative to the epithelium than the lower germ since early cap stage. Tooth engineering studies suggest that this higher proportion may help to form the supplementary cusps. Indeed, in artificial teeth made by reassociating a varying amount of mesenchymal cells to a single epithelium, the number of cusps formed increases with the number of mesenchymal cells (Hu et al., 2006). To control whether this higher mesenchyme:epithelium ratio is specific to the mouse, we extracted mesenchyme and epithelium-specific marker genes from tissue-specific transcriptomes (Figure 1C), and used *in silico* deconvolution to estimate the mesenchyme proportions from whole tooth germ transcriptomes (Figure 4A). The proportion of mesenchyme was indeed significantly higher in the upper molar in mouse, but not in hamster (Wilcoxon tests, p < 2e-16 and p = 0.152). As shown in a previous publication (Pantalacci et al., 2017), and suggested above, this change in tissue proportion should inflate the transcriptomic differences between mouse and hamster.

**Fig 4.**
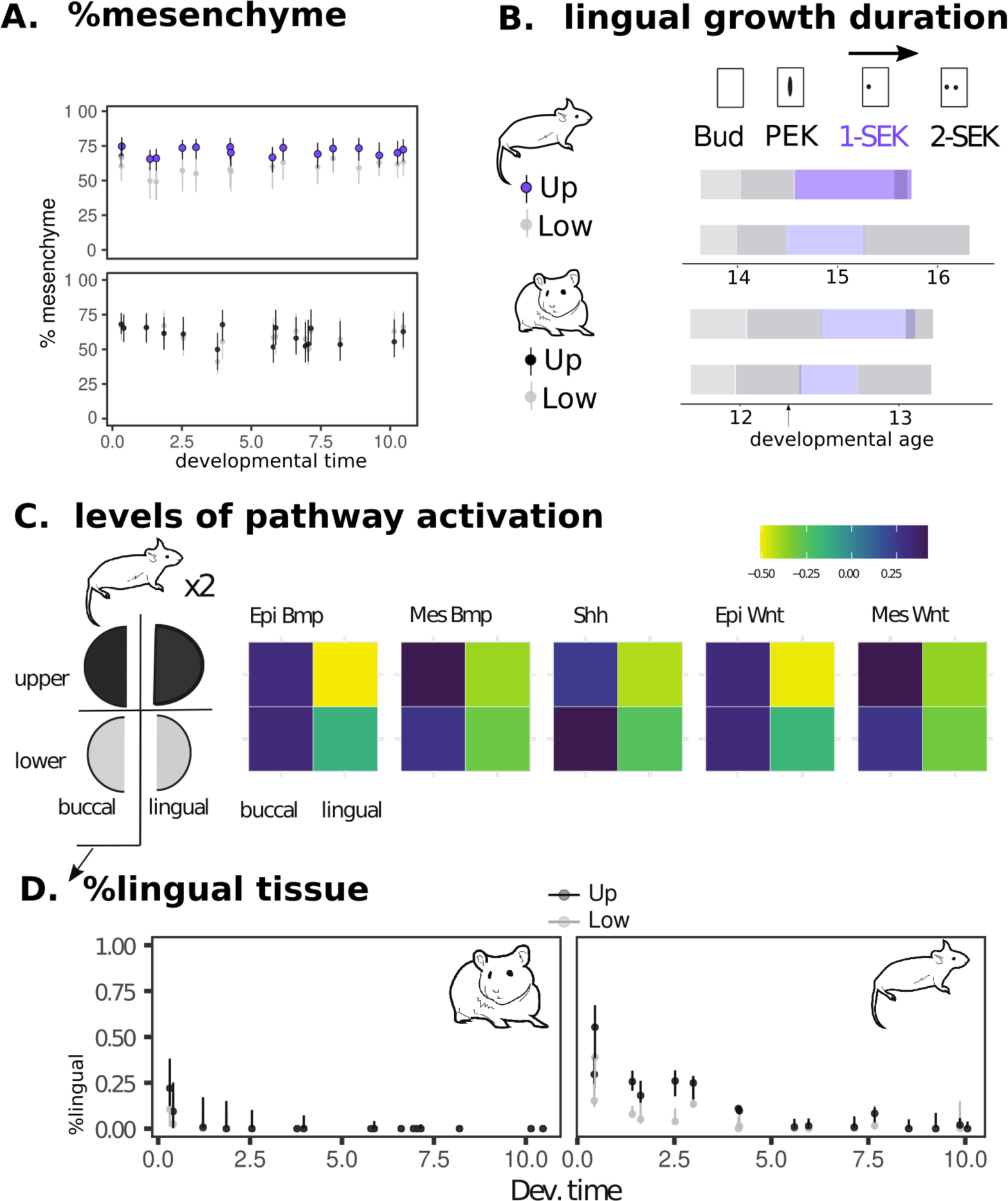
Morphogenetic changes specific to the mouse upper molar. A. Percentage of mesenchymal tissue in tooth germs estimated by deconvolution of the RNAseq time series with tissue-specific marker genes. Time scale in relative 0-10 scale (x axis) for each species. Color code indicated for each molar. B. Relative duration of the first morphological stages highlights a longer period of lingual growth in mouse upper molar (1-SEK, purple). Stage durations from Figure 1B. PEK and SEK: primary and secondary enamel knots respectively.C. Levels of activation of BMP, SHH and WNT pathways in buccal and lingual sides of the mouse molars at 15.0 dpc (1 SEK stage). Measurements made with an *in silico* method, ROMA, which compares pathway activity in transcriptomic samples based on a list of targets for the pathway. Two separate lists of target genes to estimate both an epithelial (epi) and a mesenchymal (mes) pathway activity in the 15.0 buccal and lingual RNAseq samples. Drawing on the left represents the dataset design. D. Proportion of lingual tissue in mouse and hamster molars estimated by deconvolution of the RNAseq time series with lingual and buccal marker genes.

#### A bucco-lingual polarity is maintained in mouse molars during the first steps of cusp formation, and further enhanced in the upper molar

The supplementary cusps of the upper molar form last, on the lingual side of the tooth (Figure 1B), but the dynamics of SEK formation differs much earlier between mouse and hamster upper molars. Indeed, the 1-SEK stage is longer in the mouse upper molar than in any other tooth (likelihood ratio test, p < 1e-16, Figure 4B). At that stage, the tooth germ grows rapidly on the lingual side. This finding prompted us to look into changes of the bucco-lingual development in the mouse upper molar.

We do not know the molecular mechanisms whereby the first cusp forms on the buccal side and a second one forms on its lingual side. However, we know the mechanisms which decide if a first tooth is formed, and whether a second tooth can form lingually (Jia et al., 2013, 2016; Lan et al., 2014; Zhang et al., 2009). To form a tooth, the WNT pathway must be activated on the buccal side of the mouse jaw, while on the lingual side, WNT pathway activation is prevented by OSR2, notably through the WNT inhibitor SFRP2 (Jia et al., 2016). This bucco-lingual polarity of the jaw is set up by a mutual antagonism between BMP4 activity on the buccal side and OSR2 activity on the lingual side. In mice, displacing this BMP4/OSR2 balance can suppress tooth formation (following loss of BMP4) or induce the formation of a supplementary tooth on the lingual side (following the loss of OSR2 or of the Wnt inhibitors) (Jia et al., 2013, 2016; Lan et al., 2014; Zhang et al., 2009)). Since the mechanisms for tooth (PEK) formation are largely re-used for cusps (SEK) (Jernvall & Thesleff, 2012), it would be unsurprising that this BMP4/OSR2 balance also controls the B/L polarity of cusp formation.

To look for evidence of persistent molecular B/L polarity during cusp morphogenesis, we collected and analyzed mouse transcriptomes of buccal and lingual halves at the early 1-SEK stage (Figure 1C). We found that *Osr2* and *Sfrp2* are still expressed with a strong lingual bias at 1-SEK stage, and in the timeseries, their expression is maintained at high levels during the period of bucco-lingual development of the tooth germ (Figure S7). To determine if the tooth germ is polarized at the 1-SEK stage, we estimated the levels of activation of 3 pathways controlling cusp formation (BMP, WNT and SHH), in the buccal and lingual halves separately. We used ROMA, a method that exploits the level of expression of up and down-regulated transcriptional targets to quantify signaling pathway activity directly from the transcriptomes (Martignetti et al., 2016). The list of WNT and BMP target genes was specifically established in tooth epithelium and mesenchyme by others (O’Connell et al. 2012). WNT, BMP4 and SHH pathways are strongly activated on the buccal halves of both molars, but very weakly on the lingual halves, which therefore still appears as a naive tissue (Figure 4C). In the upper molar, the lingual half looked even more naive than in the lower molar, with even lower levels of pathway activities, consistent with twice higher levels of *Sfrp2* expression, and delayed downregulation (Figures 4C, S2, S3). In the buccal half, activation of the BMP4 and WNT pathways in the mesenchyme is stronger in the upper molar, which thus appears as more polarized than the lower molar. These findings support the idea that the BMP4/OSR2 antagonism is still acting during early mouse molar morphogenesis to set up the B/L polarity of the tooth. This polarity maintains naive tissue on the lingual side of the germ at 15.0, which grows faster and shows delayed cusp formation relative to the buccal side.

We reasoned that the larger the proportion of this naive lingual tissue, the stronger the germ growth potential and its capacity to form cusps on its lingual side. We therefore quantified this proportion in mouse and hamster tooth germs, by deconvoluting the timeseries dataset with buccal and lingual tissue transcriptomes (Gong & Szustakowski, 2013). As expected due to progressive cusp formation, we found that the proportion of naive lingual tissue decreases during morphogenesis in both species (Figure 4D). But in mouse molars, and even more markedly in the mouse upper molar, the initial proportion of naive tissue is larger, and diminishes more slowly. We noted that the naive tissue proportion correlates with levels and temporal pattern of *Sfrp2* expression in mouse and hamster molars (Figure S7).

In summary, a B/L polarity of the tooth germ is maintained during the first steps of cusp formation. This is especially true in mouse molars whose proportion of lingual naive tissue is increased, and correlates with higher levels of *Sfrp2* expression. This polarization is further exaggerated in the mouse upper molar.

#### A change in the BMP pathway may underlie the maintenance of the B/L polarity in the mouse molars

The findings above suggest a change in the BMP4/OSR2 balance in mouse molars, with an exaggeration in the mouse upper molar. We came back to the list of genes with mouse upper specific profiles (Figure 3B) and compared it with the list of genes with a marked bucco-lingual bias. The *Bmper* gene is a good candidate since it is a regulator of the BMP4 pathway in many tissues, and it ranked well in both lists (respectively 211/14532 and 9/12008). In mouse early 1-SEK stage molars, *Bmper* is more expressed on the buccal side in both molars and more expressed in the upper molars on both buccal and lingual sides. In the time series, *Bmper* expression levels are higher in mouse than in hamster. It decreases with time in all teeth, but this decrease is slower in the mouse upper molar (Figure 5A). This is similar to the levels and dynamics observed for the proportion of naive lingual tissue, or the expression of *Sfrp2* gene (Figures 2C, S2). This pattern suggested us that *Bmper* participates in the regulation of the B/L polarity.

**Fig 5.**
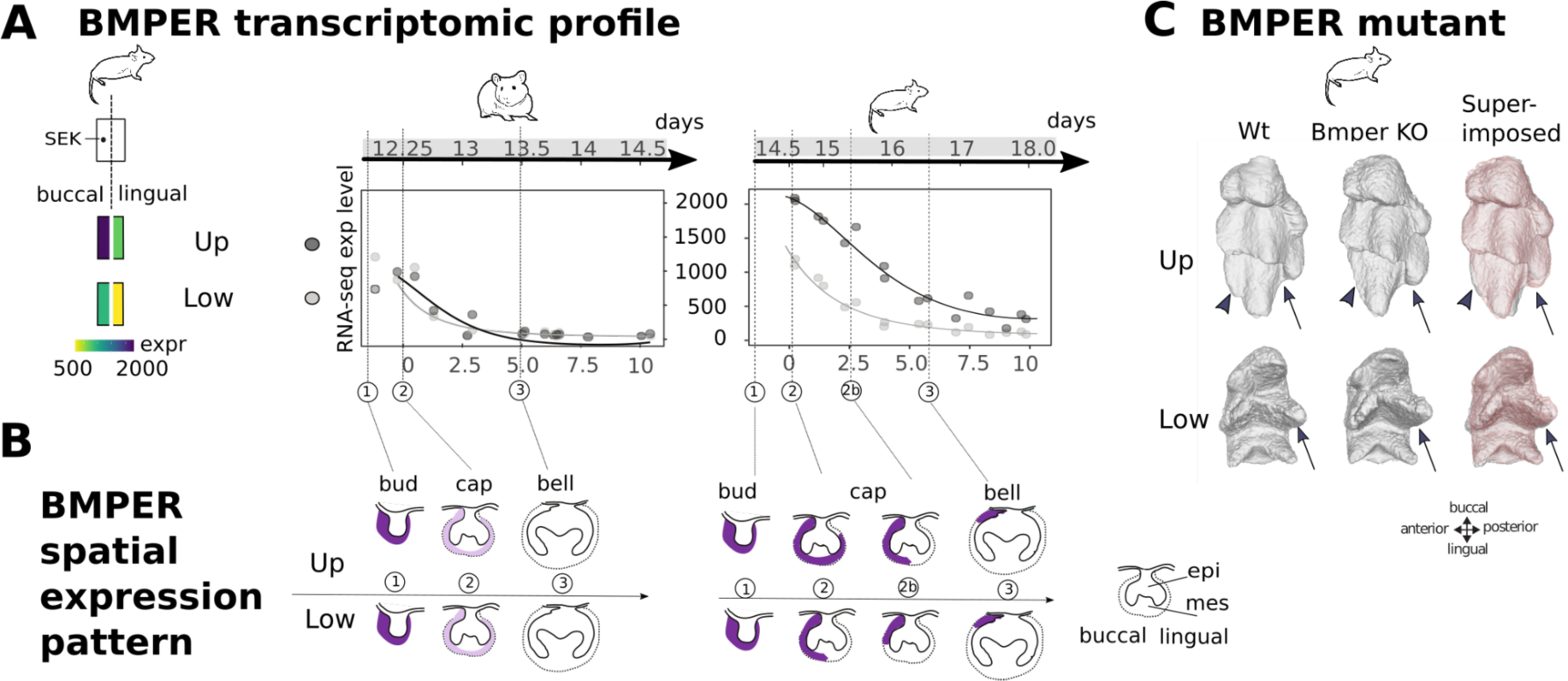
Expression and null mutant phenotype of the *Bmper* gene. A. *Bmper* transcriptomic profile. left: Colors represent expression levels in buccal and lingual parts of the mouse tooth germs (15 days). right: *Bmper* expression decreases with time and reaches a minimum when the tooth germ stops expanding on the lingual side (bell stage, n°3). Expression is higher and lasts longer in the mouse upper molar. Ages in days above the plots, in relative time on the x axis. B. Drawings of tooth germ sections at bud, cap and bell stage summarise *Bmper* expression established by *in situ* hybridization (Figure S6). Stage correspondence with dashed lines and numbers. C. Foetal molar morphology of wild type (Wt) and *Bmper* null mutant (KO). Semi-automatic reconstruction of tooth mesenchyme was performed on micro-CT scans of PTA-stained heads taken at 19.5 days. Arrows point to the enlarged lingual cusp in both mutant molars. Arrowheads point to the third-forming buccal cusp, missing in the mutant upper molar.

*In situ* hybridizations showed that in mouse molars, *Bmper* rapidly withdraws from the lingual side to remain strongly expressed in a small buccal domain only. This process is delayed in the upper molar. In the hamster, the withdrawal is symmetrical and similar in both teeth (Figures 5B, S6). Thus *Bmper* expression changed at two levels. First, it acquired a new bucco-lingual regulation leading to a strong buccal expression and an early withdrawal from the lingual side. This co-evolved in both teeth. Second, it acquired a new lower-upper molar difference, with delayed withdrawal in the upper molar.

To gain insight into the function of *Bmper*, we obtained a mouse null mutant and studied the shape of its molars. Since the homozygous *Bmper* mutant are lethal at birth, this had to be done by reconstructing the enamel-dentin junction at 19.5 days, carefully matching them with controls of similar developmental age (see material and methods). The upper molar is modified: one of the buccal cusp is absent or poorly grown (Figure 5C, arrowheads) and one lingual supplementary cusp is more prominent (arrows). The lower molar is also modified: the central lingual cusp, which is determined by formation of the 2-SEK following lingual growth, is more prominent. Thus, the loss of *Bmper* modifies the bucco-lingual equilibrium, favoring the lingual side of the molar at the expense of the buccal side. Since Bmper modulates the BMP4 pathway, this mutant phenotype reinforces the idea that the BMP4 pathway regulates B/L polarity during cusp formation, and that Bmper might have a causative role in displacing the BMP4/OSR2 balance in mouse.

In summary, we show that the BMP4/OSR2 antagonism persists in mouse to regulate the B/L polarity of the tooth during morphogenesis. As compared to hamster upper molar, the mouse upper molar has an asymmetrical expression of *Bmper* (buccal side) and an increased expression of *Sfrp2* (lingual side). This is associated with an increased and persistent proportion of naïve lingual tissue. These differences seem very consistent with the newly evolved lingual cusps of this tooth. We were very intrigued however that qualitatively, these morphogenetic changes are also seen in the lower molar, although to a lesser extent and without change in number and respective size of buccal/lingual cusps. We next wanted to quantify this concerted developmental evolution of the lower molar with the upper molar.

### Concerted evolution of lower molar development with upper molar development

#### Developmental gene expression in the lower molar largely evolve in a concerted manner with the upper molar

We modelled the temporal profiles of the 4 molars altogether to quantify their concerted evolution. We fitted four models (Figure 6A): The most complex model has four curves (one distinct per tooth), intermediate models have two curves (distinguishing species: hamster/mouse or distinguishing tooth: upper/lower), and the most simple model has a single curve common to all teeth (1 curve). Models for different species account for different baseline expression levels, to make sure that we focus on species differences in temporal dynamics. We attributed the best model to each gene and from this we built an index of coevolution. We estimated that the expression profiles of 61% genes have coevolved. This is consistent with a cruder estimate from the PCA analysis, where the main axis of variation in the transcriptomes separates samples by species, but groups upper and lower molars (Figure 1D).

**Fig 6.**
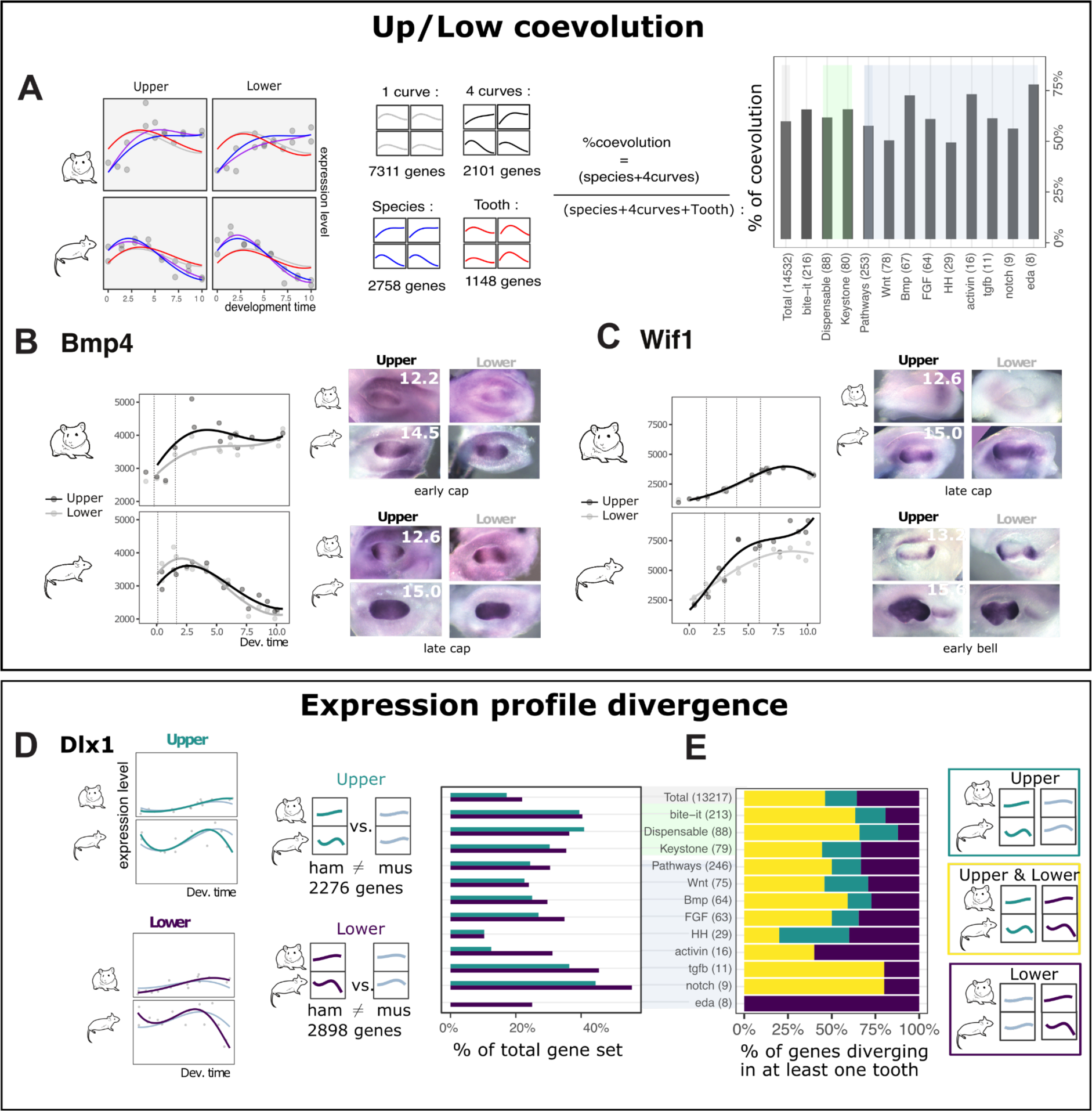
Divergence and coevolution of expression profiles. A. Nested models of temporal profiles taking the four teeth altogether. The percentage of coevolution is computed as the proportion of “divergent” genes, among genes varying between species and/or teeth. B-C. *Bmp4* and *Wif1*’s expression profiles corroborated by *in situ* hybridization of dental mesenchyme show that expression in upper and lower molar has coevolved. Dashed lines and numbers map pictures to the timeseries. See Figure S9 for details. D. Temporal profiles modelled by tooth type. *Dlx1* is shown with samples (grey dots), and models (curves). Top: the “upper divergent” model, allowing different profiles in mouse and hamster (green), is compared with the “upper non-divergent” fitting the same profile but different baseline expression levels (grey). Bottom: Same modellings fitted independently for lower molars (purple and grey). Best model was chosen for each molar by likelihood ratio test (adjusted p < 0.05). Barplots: percentage of divergent profiles in upper and in lower molars for different gene categories taken from (O’Connell et al. 2012) and (Hallikas et al., 2021). A-C, Gene categories as in Figure 2A with numbers into brackets, genes from developmental “pathways” further splitted. B. Percentage of “divergent” genes in upper and in lower molars (yellow), only in upper (green) or only in lower (purple).

The profiles of many genes important for tooth development have coevolved, suggesting that developmental processes have largely evolved in a concerted manner in the two teeth, as seen earlier for the B/L axis. We decided to examine by *in situ* hybridization some of those genes whose profile co-evolved in the transcriptomes. We wished to determine what spatio-temporal profiles are behind this transcriptomic co-evolution, and which other developmental processes besides B/L polarity may have co-evolved between the two teeth.

#### The early dynamics of cusp formation show concerted evolution, with anticipated cusp formation in mouse

We first examined Bmp4, since finding this essential gene for tooth development among co-evolving genes was a surprise (Figure 6B). *Bmp4* is expressed in both the epithelium and the mesenchyme, but the epithelial domain is so small relative to the mesenchymal domain (see later Figure 7 and S9), that the latter will dictate the bulk transcriptomic profile. We therefore first looked at the *Bmp4* expression in the mesenchyme. *Bmp4* reached a spatially homogeneous expression earlier in the mouse mesenchyme (Figure 6B), consistent with an earlier peak of expression in mouse transcriptomes.

**Fig 7.**
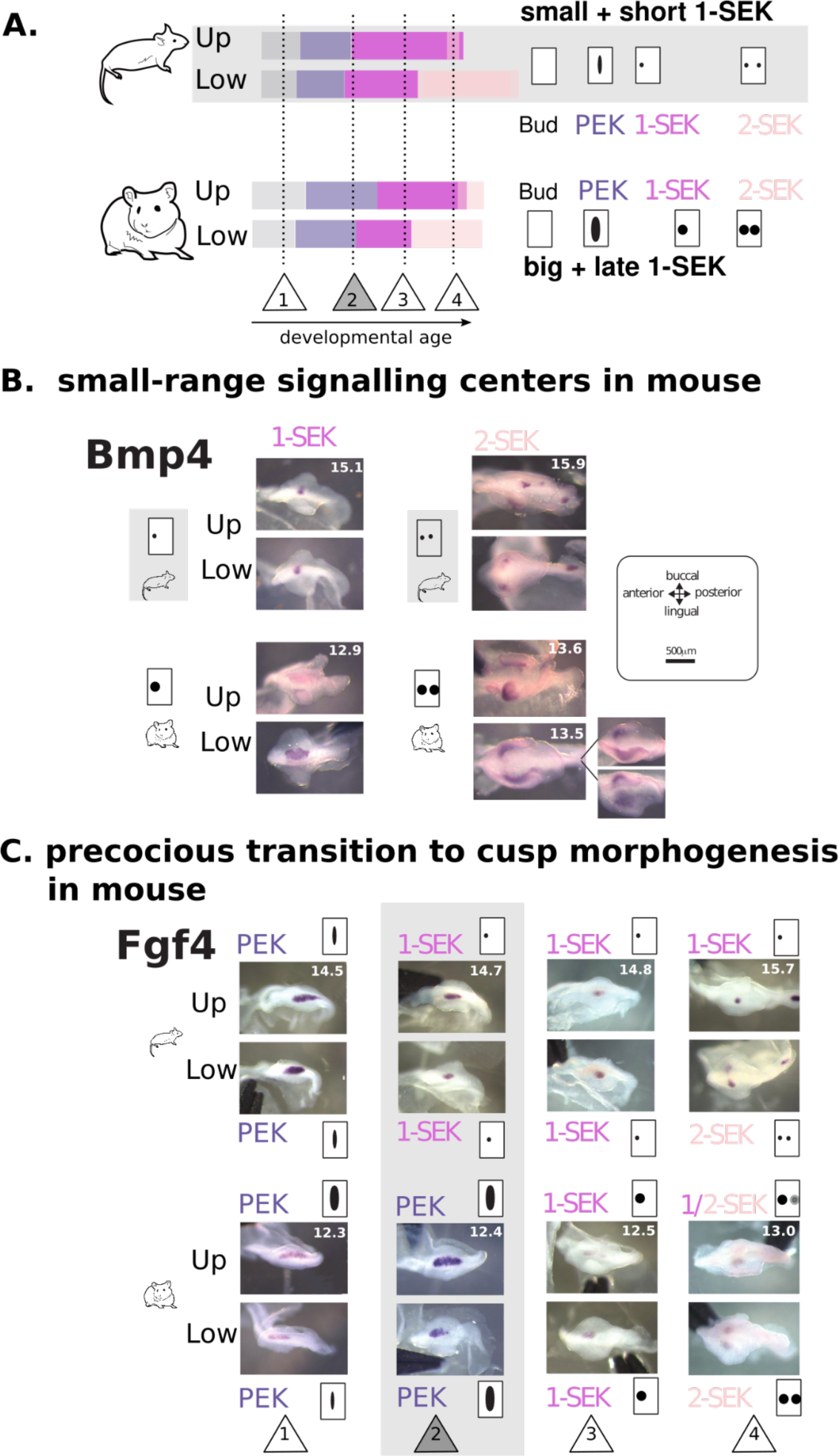
Activation-inhibition mechanisms co-evolved in mouse molars. A. Duration of each stage estimated by Markov models from Figure 1B. Numbers in triangles as in B. Timeline and cartoons on the right recapitulate changes in signalling centres. B. Mouse molars transition earlier to cusp patterning. Transition from the PEK to the 2-SEK stage as seen on tooth germ epithelial parts hybridised against *Fgf4*. Pairs of mouse/hamster embryos were selected to show four remarkable steps in this chronology (1-4 in triangles). At stage 2, *Fgf4* expression is still elongated in hamster, as typical for a PEK, while it is already roundish in mouse, as typical for a SEK. C. Expression of *Bmp4* is more focalized in mouse than in hamster SEKs. Mouse and hamster samples are paired for similar advancement of epithelial growth. Age of samples in days is in the upper molar picture (for samples taken from the same embryo) or in both pictures (samples taken from different embryos). See also Figure S10.

We picked up two other mesenchymally expressed genes for in situ hybridization, Wif1 and Dkk1, because 1) these genes are likely involved in the gene regulatory network of tooth formation: they are known modulators of the Wnt pathway which is critical for cusp formation, and their expression is changed when the Bmp4 and/or Wnt pathways are manipulated (data by O’Connell et al.) and 2) their temporal profiles markedly differ from each other, as well as from *Bmp4*.

*Wif1* rises earlier in mouse transcriptomes, and its mesenchymal expression is seen both earlier and in a larger territory in mouse tooth germs (Figures 6C, S9). *Dkk1* expression transiently decreases in mouse transcriptomes, which coincides with an earlier relocalisation of its expression at future cusp tips beneath the SEK (Figure S9). This expression pattern suggested to us that cusp formation might in fact be anticipated in mouse as compared to hamster.

We thus turned back to our quantification of cusp formation dynamics, and realized that both mouse molars quickly transition to 1-SEK after a rather short PEK stage (Figures 1, 7A). By comparing epitheliums of the two species matched for growth advancement, we found that both mouse molars already exhibit the rounded and focalized *Fgf4* expression typical of a SEK when hamster’s still exhibit the large and elongated *Fgf4* expression typical of the PEK (Figure 7B stage 2). Thus, mouse PEK is rapidly turned into a precocious and focalized 1-SEK, marking an early beginning of cusp patterning in a very young cap stage, and this happens very similarly for both mouse molars. The dynamics of activation-inhibition mechanisms relative to the advancement of epithelial growth thus evolved in a concerted manner in the two teeth. We have shown previously how the transcriptome carries signatures of ongoing developmental processes, including cusp formation (Pantalacci et al., 2017). We suspect that well beyond the few genes studied above, the transcriptome could be deeply shaped by this anticipated formation of cusps in the young mouse molars, and thereby be markedly different from hamster transcriptomes. Said differently, this morphogenetic change could contribute to a substantial part of the transcriptomic concerted evolution.

#### Spatial aspects of the activation-inhibition mechanisms controlling cusp formation also show concerted evolution, with more local inhibition in mouse

To get further insight into the evolution of activation-inhibition mechanisms in hamster and mouse, we next studied the expression of diffusing signals, which are produced in the SEK and inhibit the formation of other SEKs in the vicinity. Indeed, evolutionary changes in the production and diffusion of these signals are thought to drive evolutionary changes in cusp number (Salazar-Ciudad & Jernvall, 2010). We do not know how much each of these molecules diffuses, but the spatial range of expression of each gene can be considered as a minimal range for its inhibitory action on cusp formation. We studied two known inhibitors of SEK formation, *Bmp4* (Meguro et al., 2019) and *Shh* (Kim et al., 2019). We also studied *Wif1*, which we consider a likely inhibitor since it antagonizes the Wnt pathway, whose activation in the epithelium promotes cusp formation (Järvinen et al., 2006; Liu et al., 2008). *Bmp4* and *Wif1* are expressed from the signalling centers : their expression pattern is much more narrow and roundish in mouse than in hamster (both in PEK and SEKs) (Figures 7B and S10). *Shh,* which more largely marks cells committed to cusp formation, also shows a more restricted expression in mouse (Figure S10). Hence for the three pathways involved in the activation-inhibition mechanisms, inhibition is more local in both mouse molars.

In conclusion, both dynamics and spatiality of activation-inhibition mechanisms have evolved in a concerted manner in the molars of the two species. Both of them, rapid switch to SEK formation relative to epithelial growth and a more local inhibition, are predicted to favor the formation of more cusps. Features that make sense for the formation of the supplementary cusps in the upper molar are thus observed also in the lower molar, as seen above for the B/L polarity.

### Developmental gene expression diverged as much in lower as in upper molar

These developmental phenotypes of the lower molar have evolved in concert with the upper molar, but this evolution did not drive a major phenotypic change. Such discrepancy between the divergence of development and the conservation of the final phenotype, is a phenomenon known as Developmental System Drift (DSD). To measure the extent of this phenomenon, we decided to compare levels of developmental evolution in both teeth.

Since the lower molar phenotype has been much more conserved during evolution, the lower molar developmental phenotype captured by the temporal profiles should be more conserved. Otherwise, this is an indication of DSD.

We scored the divergence between mouse and hamster upper and lower molars by modelling temporal profiles with polynomials (LRT with adjusted p < 0.05). We found that for 22.0% of genes, the profiles diverged in the lower molar, which is even more than in the upper molar (17.3%, Figure 6D, Table S2). This is true as well for genes relevant for tooth development and phenotype (“bite-it”, “keystone”, “pathways”; Figure 6E). Put together, these observations suggest that the development of the lower molar has drifted while co-evolving with the upper molar.

### Bat limbs development show a similar pattern of concerted evolution

In order to generalize our results, we turned to another case of drastic independent evolution: the bat limb. The evolution of the wing relied on drastic changes in the forelimb development, including changes in digit patterning, growth, and webbing to form the wing membrane. In comparison, the bat hindlimb kept a morphology more typical of quadrupedal species, as did both mouse limbs. This provides a framework of independent evolution allowing us to test the generality of our findings beyond mouse molars.

We collected raw sequencing data from a previous study comprising 3 stages of mouse and bat fore/hindlimb development (Maier et al. 2017). We quantified expression levels and classified temporal profiles with polynomial models dedicated to measure coevolution (as in Figure 6A).

Just like in our molar dataset and consistent with the original analysis of this dataset (Maier et al., 2017), the genes which have a limb-specific temporal profile and which have kept it in mouse and bat are a minority (53 genes), but they are highly enriched for transcription factors, including all the expected identity genes, eg. *Tbx4* and *Tbx5*, *Pitx1*, as well as 6 biased Hox genes.

The profiles of 714 genes differed both between species and limbs. The profiles of almost four times more genes (2677) diverged between the two species, but co-evolved in the two limbs, despite their drastic morphological differences. Such a large proportion of co-evolving genes mirrors our finding in rodent molars. Importantly, genes with a well-established role in controlling limb morphology co-evolved. It is the case of key genes controlling limb patterning (*Shh*, *Fgf10*, *Fgf8*, *Grem1*…) and chondrogenesis (*Wnt3* and the Activin pathway: *Inhba*, *Inhbb*, *Acvr2b*…). It is also the case of most of the genes known to regulate webbing (*Fgf8*, *Grem1*, *Bmp7*, *Ihh,* Retinoic acid pathway: *Aldh1a2*, *Cyp26b1*).

We note that several of these genes have been pointed in the literature as key for bat wing evolution. For three of them, we could compare the expression profile in the transcriptomic dataset with published *in situ* hybridization in both limbs, and they were consistent with co-evolution. The iconic *Shh* gene expression clearly peaks at the second stage in both bat limbs, but not in mouse limbs (Figure S11), and peaking is exaggerated in the bat forelimb (Figure S11). This is consistent with figure 2 in (Hockman et al., 2008). The new temporal profiles of *Fgf8* and *Grem1* in both bat limbs are also consistent with a previous study, which has shown the novel expression domain of these genes in both limbs (Weatherbee et al., 2006).

As in mouse molars, co-evolution is pervasive in bats limbs and concerns genes whose expression evolution was key for the independent phenotypic evolution of the forelimb.

## DISCUSSION

Below we discuss how the independent phenotypic evolution of the mouse upper molar involved reinforcing and building on ancestral specificities of the upper molar development, in relation with identity genes. It was accompanied by extensive evolution of lower molar development, including concerted evolution with upper molar development, which contrasts with the limited phenotypic evolution in this tooth. These findings are best understood in a model where developmental system evolution of the upper molar induced developmental system drift in the lower molar.

### Conserved specificities of lower and upper molar morphogenesis may date back to early mammals

We found several conserved specificities which discriminate between lower and upper molars. All mark the early period of cusp formation: the arrangement of cusps at 3-SEK stage and the early dynamics of cusp formation, the morphology of lingual epithelium and a transient reinforcement of transcriptomic identity (already seen in mouse in our previous study (Pantalacci et al., 2017)). Since these findings were made in hamster and mouse, these specificities of lower and upper molar were likely present in their common ancestor, but we suspected they may even date from early mammals.

Early mammals evolved “tribosphenic molars”, a major innovation of lower and upper molar shape which enabled unprecedented occlusion (B. M. Davis, 2011; Hillson, 2005). For the first time in the reptilian evolution, lower and upper molars were developing into drastically different shapes. In the Figure S5, we discuss in detail how the developmental specificities of lower and upper molars of mouse and hamster strikingly mirror the specificities of the lower and upper tribosphenic molars, taking into consideration known homologies in mammalian molar cusps. In particular, we show how the spatio-temporal dynamics of cusp formation in the lingual and posterior directions are combined differently in the two molars. We propose that evolving a jaw-specific control of this combination was the key developmental innovation underlying the invention of the tribosphenic molars.

Most later mammals had less dissimilar teeth, such as the common ancestor of mouse and hamster or the present golden hamster. Yet the heritage of the mammalian innovation remains visible in the transient developmental dynamics of lower and upper molars. This also constitutes a case of “recapitulation” since early ontogeny of cusp formation recapitulates phylogeny (Gould, 1977).

These hidden developmental specificities could serve as a basis for the independent evolution of upper molar in the mouse lineage. It is interesting that some fossil rodents close to mouse/hamster common ancestor carried a crest along the lingual basis of the upper, but not the lower molar, which may be seen as an ancestral predisposition to enlarge lingually and form lingual cusps (Charles et al., 2009; Tiphaine et al., 2013).

### What does underlie the conserved morphogenetic identity of molars? Ancestral molecular identity of molars

We identified several conserved specificities of upper and lower molars at the developmental system level, yet such conserved specificities remain rather discrete at the transcriptomic level. We found relatively few genes consistently biased in the two species, and their temporal profiles were most often not conserved, except some highly relevant transcription factors. This includes the two jaw-specific genes *Nkx2-3* and *Pou3f3,* and several *Dlx* genes. The *Dlx* genes are homeobox transcription factors which specify jaw identity at early stages of craniofacial development in jawed vertebrates and might have been implicated in the transition from a reptilian to a mammalian jaw (Depew et al., 2005; Gillis et al., 2013). The dose as well as the complement of *Dlx* genes (*Dlx1/2* in upper jaw; *Dlx1/2/5/6* in the lower jaw) are important for normal jaw development in mouse (Depew et al., 2005), and upper molars fail to develop without *Dlx1/2 (Qiu et al., 1997)*. *Dlx1*/2 showed an upper-bias in both species, and Dlx5/6 a lower bias. Transcription factors known to be essential to tooth development, such as *Msx1*, *Barx1*, *Pitx1* also showed a conserved bias. It remains to be tested if this bias is directly controlled by identity genes.

Our results show that during evolution, the details of developmental interactions in serial organs diverge extensively, but some developmental specificities of one organ with respect to the other are conserved (e.g. the relative order and timing of appearance of the 3 first cusps and the period of maximal transcriptomic divergence, the delayed development of the upper molar). These specificities could be encoded in a conserved relative dose of the key transcription factors specifying an organ (here a molar or a limb), and this conserved relative dose could be controlled by identity genes. Altogether this forms an ancestral molecular and developmental identity for the two teeth.

### Reinforcement of the ancestral molecular and morphogenetic identity in mouse molars We found that the molecular identity was reinforced in mouse, in the upper molar but also

more surprisingly in lower molar. Expression levels doubled for the upper-molar specific TF *Pou3f3* and the lower-molar specific TF *Nkx2-3*, and the ancestral bias of many genes was exaggerated, whether in favor of upper or lower molar (e.g. *Barx1* and *Dlx1*; *Pitx1*, respectively). Consistent with these changes in individual TFs, we found at genome-wide scale that the temporal profiles in the mouse were exaggeratedly different from each other. As for specific TF, divergence is seen both in upper and lower molar.

### Mouse upper molar displays three morphogenetic changes favoring supplementary cusp formation, one is building on ancestral upper molar specificities

The supplementary cusps form last, the most anterior one being the very last, just as it happens in the fossil record (Lazzari et al., 2008; Tiphaine et al., 2013). This “recapitulation” looks superficially like a case of “terminal addition”, a mechanism for evolutionary change whereby development is incremented with one more step (Gould, 1977). However, species divergence peaks early in development and we point to three features that concern mouse early upper molar development, but could pave the way for the supplementary cusps (Figure 8).

**Fig 8.**
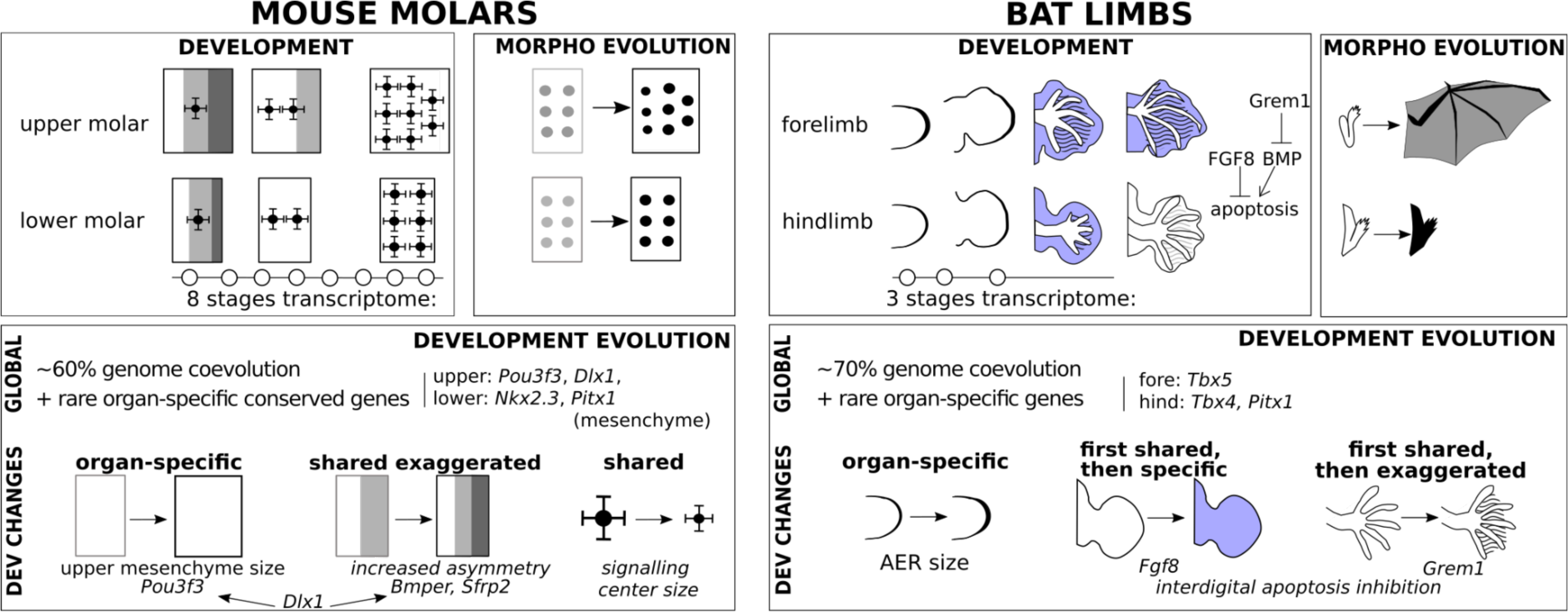
A summary of findings and working model from this study. For mouse molars and bat limbs, the “development” box shows key developmental stages with the time period covered by transcriptome data. The box “morphological evolution” represents the drastic morphological changes of the upper molar and forelimb as compared to the relative conservation of the lower molar and hindlimb. The “development evolution” box summarises the evolution of transcriptome and developmental mechanisms. Transcriptomes are dominated by co-evolution, with rare conservation of organ-specific expression besides identity genes. Organ-specific, shared, and shared but exaggerated developmental changes combine to achieve the organ-specific morphological change. For each change, the ancestral and derived state are represented, and candidate genes are indicated. In mouse molars, 3 changes (increased mesenchyme size, increased bucco-lingual asymmetry and smaller inhibitory signalling centres) combine to induce extra cusps on the lingual side of the upper molar only. In bat limbs, we took the example of early changes in AER size (apical ectodermal ridge, a signalling center) leading to altered digit patterning and late expression changes in *Fgf8* and *Grem1*, that efficiently combine to suppress interdigital apoptosis in the forelimb only. In our working model, shared changes are necessary but not sufficient. Combining them with more specific changes (e.g. mesenchyme size) and/or ancestral specific features (e.g. ancestral difference in molar Bucco/lingual axis) is necessary to achieve the morphological change. Related to Figure S12-13.

The first feature is the larger mesenchyme compartment of the mouse upper molar. Increasing mesenchyme proportion increases cusp number in tooth engineering studies (Hu et al., 2006), probably because the mesenchyme promotes epithelial growth, which enlarges the field where activation-inhibition mechanisms act and pattern SEKs. The observed difference seems however too modest to drive the formation of supplementary cusps on its own and does not explain why the supplementary cusps would form on the lingual side only.

The second feature is the stronger polarisation of the upper molar field along the bucco-lingual axis, associated with a precocious transition from PEK to 1-SEK and a very long 1-SEK stage. As a consequence, a larger undifferentiated field is present on the lingual side, where activation-inhibition loops can pattern SEKs. This feature seems highly relevant because it could explain why the increase in cusp number is focused on the lingual side of the tooth, while the size of buccal cusps is reduced. It seems to exploit an ancestral specificity of the upper molar as compared to the lower molar, which produces a longer 1-SEK stage. This specificity could combine together with a novelty in mouse responsible for shortening the PEK stage in both teeth, and produce an upper molar with a very long 1-SEK stage and a large undifferentiated field, while change remains more modest in the lower molar.

The third feature is the narrower range of expression of signaling molecules in mouse signalling centres, which is especially obvious for *Bmp4*. Reducing the range of these known cusp formation inhibitors should allow to squeeze more SEKs in an equivalent field. In agreement with this idea, a mouse mutant where *Bmp4* is overexpressed in all the epithelium loses the supplementary cusps (Meguro et al., 2019), hence reverting to the ancestral phenotype.

### Molecular mechanisms and candidate genes for the observed morphogenetic changes

Mapping mutations corresponding to these developmental phenotypes is out of the scope of this study, but transcriptomics provided us with molecular mechanisms and some candidate genes. No expression change was observed in two obvious candidates from the literature (*Fgf3* and *Activin*⍰*A*, Figure S12, (Charles et al., 2009; Kwon et al., 2017).

In insects, the evolution of the dose of identity genes has been correlated with the evolution of the size of serial organs (Paul et al., 2021). *Pou3f3* is expressed specifically in the upper molar mesenchyme of both species and its dose is twice increased in mouse. Hence *Pou3f3* is a good candidate to explain the larger proportion of mesenchyme specifically in the mouse upper molar. *Pou3f3* loss-of-function mutants miss some skeletal elements of the upper jaw, but their upper molars showed “no major defects” (data unshown in (Jeong et al., 2008)). This may deserve re-examination, or study in sensitized backgrounds. The causative mutation may also be upstream in the regulatory network, in particular in *Dlx1/2*, since *Pou3f3* is regulated by *Dlx1*/2 in the early jaw (Jeong et al., 2008) and the dose of *Dlx1/2* genes are also twofold increased. Both *Dlx1/2* and *Pou3f3* are also involved in cranio-facial development. Since mastication has changed together with tooth morphology in the mouse lineage (Lazzari et al., 2008; Tiphaine et al., 2013), changes in the dose of these genes could have had pleiotropic effects beyond the molar.

The reinforcement of the B/L polarity in the mouse upper molar likely involved changes in or upstream of the BMP4/OSR2 network. Interestingly, mutations in this network have very different consequences on lower and upper molar development of mouse (eg. a tooth is normally formed *versus* arrested at a very early stage) (Jia et al., 2013, 2016; Kwon et al., 2017; Lan et al., 2014). We show that the BMP4/OSR2 antagonism, which earlier in development regulates the B/L polarization of the molar-forming region, persists in mouse during the first steps of cusp formation. We found expression changes in two genes which should modify the output of this BMP4/Ors2 network: *Sfrp2*, whose role in the network was known in mouse (Jia et al., 2013, 2016; Kwon et al., 2017; Lan et al., 2014) and *Bmper*, whose role in tooth development was unknown.

BMPER is a known modulator of the BMP4 pathway, which seems to be pro- or anti-BMP4 in different contexts (Correns et al., 2021; Ikeya et al., 2010; Kelley et al., 2009; Serpe et al., 2008). Because the molars of the *Bmper* loss-of-function mutant are “hypo-buccal/hyper-lingual”, we deduce that *Bmper* is normally pro-BMP4 in the mouse BMP4/OSR2 balance. In mouse, the strong buccal *Bmper* expression should favor BMP4 activity on the buccal side while the early lingual withdrawal should decrease BMP4 activity on the lingual side. This sharper gradient of BMP4/OSR2 antagonism may link two observations in mouse: on the buccal side, the earlier PEK/SEK transition; on the lingual side, the larger undifferentiated field. Finally, we note that the teeth of the *Bmper* mutant are strikingly similar to the molars of a mouse relative, *Mastacomys fuscus brazenori*. Its first upper molars have very large supplementary cusps and have lost the same buccal cusp as the *Bmper* mutant, and its lower molars have larger lingual cusps as compared to buccal cusps (Museums Victoria Collections). Therefore, our data strongly suggest that evolutionary changes in the BMP4/OR2 network might be responsible for both the evolution of the murine dental plan and its further diversification.

Where could be the mutation which impacted this BMP4/Osr2 network? From the present data, we envision at least three possibilities. 1) A cis-regulatory change in the *Bmper* gene could have led to its new asymmetric profile, and feedback on *Sfrp2* expression through the network. In cichlids, a QTL containing *Bmper* is associated with variation in tooth number(Bloomquist et al., 2015). Species with more teeth have reduced *Bmper* expression, as mouse with more lingual cusps have reduced lingual *Bmper* expression (Bloomquist et al., 2015). 2) The mutation may lie in *Sfrp2*, and feedback on *Bmper* expression. 3) The mutation might also lie in *Dlx1/2* genes, because they control both *Sfrp2* and *Bmper* expression levels in the early mouse jaws (Jeong et al., 2008). It is striking that these 3 genes show only a minor expression difference in favor of the upper molar in hamster, but their expression is twice increased in mouse, together with a sharper upper molar bias. By acting on both a buccal (*Bmper*) and a lingual *(Sfrp2)* gene with antagonistic effects on the BMP4/Osr2 balance, the increased dose of *Dlx1/2* might have converted the mild ancestral polarization of the tooth into the sharp bucco-lingual polarisation seen in mouse. Future work focusing on the evolution of the cis-regulatory regions of these genes could test these hypotheses.

Finally, we noticed clear changes in the expression patterns of *Bmp4*, *Wif1* and *Dkk1*, three members of the BMP and Wnt pathways at the core of activation-inhibition networks (O’Connell et al., 2012; Salazar-Ciudad, 2012). Given the many regulatory feedback in these networks, a mutation may lie in one of these genes and feedback on the expression of the others, or lie in another gene to be identified.

### Lower molar development evolved in a concerted manner with upper molar development

We show that the lower molar development has coevolved with the upper molar development. Temporal expression profiles coevolved massively and several features of cusp formation also evolved in a concerted manner: the precocious PEK to SEK transition, the narrow expression of inhibitors in signaling centers, the marked bucco-lingual asymmetry with persistence of some lingual naive tissue and the early *Bmper* withdrawal from the lingual side. Concerted evolution in bucco-lingual development is especially striking, since neither cusp number nor relative size along the B/L axis differs between mouse and hamster lower molars (Figure 1). The only derived features of the mouse lower molar are the connection between cusps (the crest connecting central and posterior cusps is lost) and their slightly more parallel arrangement (Lazzari et al., 2008).

Why these concerted developmental changes translate into minor phenotypic change in the lower molar but major ones in the upper molar is an open question. Molar development has non-linear properties, characterized by threshold effects (Gjuvsland et al., 2013; Milocco & Salazar-Ciudad, 2020; Morita et al., 2020; Urdy et al., 2016). Since the lower molar can form supplementary lingual cusps when activation is boosted by adding ACTIVINβA to the culture medium (Harjunmaa et al., 2012), it suggests that in wild type mouse, the lower molar remains below a threshold, while the upper molar passes it. There could be two different, non mutually exclusive reasons for that: 1) As mentioned above, there were ancestral differences in the regulation of the bucco-lingual axis in the common ancestor. When facing the same expression change (e.g. early *Bmper* withdrawal from the lingual side), its upper and lower molar might then have reacted very differently, passing or not the threshold. 2) The upper-specific increase in mesenchyme proportion may be just the small effect needed to pass the threshold in the upper molar, while the lower molar remains below it. Indeed, the mesenchyme is the endogenous source of ACTIVIN**β**A, whose supplementation produces lingual cusps in the lower molar (Harjunmaa et al., 2012).

### A combinatory model to explain the independent phenotypic evolution of the upper molar with concerted developmental evolution in the lower molar

The three features that we observed hint at very complementary aspects of tooth development (Figure 8). The tooth literature shows it is difficult to increase cusp number in mouse molars: *in vitro* experiments have shown it can be necessary to play on multiple pathways, and mouse mutants show, at best, small accessory cusps, but no supplementary main cusps (Harjunmaa et al., 2012). None of these three changes should be sufficient on its own to induce the major changes in cusp size and proportions seen in mouse as compared to its ancestor. We therefore propose that the new phenotype involves combining mutations in at least two or three different genes, corresponding to these three features (Figure 8). Such a model with additive changes is also coherent with the stepwise addition of the supplementary cusps in the fossil record. Stem murine rodents had a single small extra cusp. Enlargement of this cusp, addition of a second extra-cusp, and size reduction of the buccal cusps came later (Tiphaine et al., 2013).

The 3 mutations which could lie behind the observed developmental phenotypes represent three different categories, with respect to their consequences for the lower molar (Figure 8).

i) mutation with organ-specific developmental effects - A mutation in the upper-molar specific *Pou3f3* gene which was part of the ancestral lower/upper code could have molecular effects specific to the upper molar. Such a mutation and effect are expected from the abundant literature on homeotic genes and serial appendage evolution.

ii) mutation with shared effects, but exaggerated in the upper molar - Another mutation could have the same molecular effects on the development of both molars (e.g. a mutation making *Bmper* expression asymmetric) but with a stronger expressivity in upper molar (e.g. larger lingual field) because the ancestral lower/upper code determines a different developmental context between the two teeth.

iii) mutation with fully shared effects on lower and upper molar development - A third mutation could have the same molecular effects and the same expressivity on the developmental phenotype (range of inhibitor expression in the signaling centers of the two teeth), but because it cannot combine with other effects in the lower molar as it does in the upper molar, it might have a very limited impact on the lower molar phenotype.

Such a genetic model is consistent with findings in butterflies’ wings. Indeed, combinatory effects of mutations and context-dependency on the ancestral homeotic code have been proposed to explain the evolution of eyespot patterns in the fore and hind-wings (Monteiro 2007, 2021, 2022).

### The patterns of transcriptome evolution seen in teeth resemble patterns observed in other serial organs

Comparative transcriptomics in embryos may be confounded by methodological effects that could inflate interspecies differences in expression levels. This includes estimating expression levels with RNA-seq data from different species and sampling a few stages in a continuous developmental window in species with different developmental rates. We controlled for this by estimating expression levels on orthologous portions of the transcripts, by matching the time window with homologous stages, by sampling many time points, and working on temporal profiles instead of individual stages.

To the best of our knowledge, there are only two other studies using interspecies transcriptomics in serial organ evolution, in similar settings: at least two species, one representing ancestral morphologies, and another one where a single organ strongly diverged from the ancestral morphology. One study compares bones from fore/hind limbs in mouse and jerboa at a single timepoint (Saxena et al., 2022). The other compares limbs/wings in mouse and bat at three timepoints (Maier et al., 2017) which allowed us to reanalyse the data with our methods.

Similar patterns are seen in all these datasets. 1) Only a small set of genes, enriched in transcription factors, showed an organ-specific expression conserved between species (our tooth data, reanalyzed limbs dataset, and (Maier et al., 2017; Saxena et al., 2022)). 2) The expression of large numbers of genes co-evolved in the two organs (tooth and reanalyzed limb data, (Saxena et al., 2022)). 3) Expression differences between serial organs are increased in species with the morphological innovation (tooth and reanalyzed limbs data, (Saxena et al., 2022)) 4) but the serial organ which kept the most ancestral morphology does not show better expression conservation (tooth and reanalyzed limbs data, and (Saxena et al., 2022).

### Co-evolution is also pervasive in bat limbs, where adaptation combines organ-specific with shared gene expression changes

Our comparative analysis of early mouse and bat development revealed that developmental dynamics of gene expression is largely shared by the two bat limbs, despite their drastically different morphologies. This concerted evolution was largely overlooked so far (e.g. *Shh* gene), because attention was mainly given to wing-specific developmental features, which seem more logically susceptible to explain wing evolution. We however realized that gene expression changes, previously pointed for their role in bat wing evolution, are in fact accompanied by concerted expression changes in the hindlimb. The bat wing membrane is achieved by suppressing the apoptosis which normally defines the digits. Functional tests showed that this is achieved by simultaneously activating the anti-apoptotic FGF pathway and downregulating the pro-apoptotic BMP pathway (Weatherbee et al., 2006). *Fgf8* and the BMP inhibitor *Grem1* coevolved in our analysis (Figure S12), with a new mesenchymal expression in both bat limbs (Figure 8, drawn from Figure 3 A-C, E-G in (Weatherbee et al., 2006)). At later stages, *Fgf8* mesenchymal expression persists in the interdigital area in the wing, but not in the foot (from Figure 3H in (Weatherbee et al., 2006)). In contrast, the BMP inhibitor *Grem1* expression persists in both limbs, with higher levels in the wing (from their Figure 3D, note the blue staining remaining around digits whereas more proximal parts of the limb are not stained at all). Thus, at this stage, specific and exaggerated shared gene expression changes seem to combine to pass the threshold for apoptosis suppression in the wing, but not in the foot (Figure 8). This evolutionary scenario of independent evolution is thus very similar to teeth, involving a combination of specific, shared, and exaggerated shared expression changes and differential threshold effects.

### Concerted evolution with Developmental System Drift is a mechanism facilitating independent evolution of serial organs

We observed incongruent patterns of transcriptome and morphologies in molar evolution : transcriptomes diverged equally in the upper and lower molars, while the morphology of the lower molar remains largely conserved. This unexpected level of developmental divergence as compared to morphological conservation is called Developmental System Drift (DSD, (Cutter & Bundus, 2020; Félix, 2012; True & Haag, 2001)).

There is now accumulating evidence that cryptic changes in developmental systems are frequent in evolution (Félix, 2007; Guignard et al., 2020; Torres Cleuren et al., 2019; Wotton et al., 2015). Because natural selection mainly acts on the final product of development, divergent developmental paths may be taken to reach the same final phenotype and drift in development is neutral with respect to natural selection. Further taking into account that genomes are constantly mutating, DSD appears as a likely alternative to developmental conservation (Félix & Wagner, 2008; Peter & Davidson, 2011).

The situation here seems different from this classical definition of DSD since at least part of lower molar and hindlimb DSD is not random: it is concerted with developmental innovation in the other organ, and therefore likely induced by the adaptation of this other organ. Because the lower molar and the hindlimb developmental systems could evolve while robustly maintaining the final phenotype during evolutionary times, mutations with shared effect could be used by adaptation. This is unexpected since it is commonly thought that adaptive mutations need to be modular at the DNA level to have organ-specific effects and thereby circumvent gene pleiotropy. The capacity of developmental systems to undergo DSD is another way of circumventing gene pleiotropy, and thus appears as a mechanism by which non-modular mutations can be selected in adaptation. We propose this is the reason why independent evolution can be so frequently seen in nature despite gene pleiotropy.

### Pleiotropy, concerted evolution and DSD

Serial organs such as molars and limbs have a heavy pleiotropy load and for this reason, they are possibly especially prone to developmental co-evolution and DSD. We nevertheless believe that our results in serial organs illustrate a much more general correlation between pleiotropy and DSD at the organismal level, as suggested previously (Félix, 2007; Pavlicev & Wagner, 2012).

The link between pleiotropy and DSD has been observed in experiments of *in silico* evolution (Johnson & Porter, 2007; Tulchinsky et al., 2014). It has also been observed in nematode genetics with a mutation increasing the fitness in laboratory conditions that has induced DSD in the vulva (Duveau & Félix, 2012). Finally, a link between pleiotropy and concerted transcriptomic evolution has already been suggested. In most multispecies transcriptomic analyses, samples of different organs tend to group by species (like molar samples in Figure 1C). This pattern, so-called “species signal”, often dominates in samples of adult tissues (e.g. kidney, brain, liver… (Brawand et al., 2011)) as well as in individual embryonic timepoints (Liang et al., 2018; Tschopp et al., 2014). This has been reinterpreted as a conspicuous concerted evolution, possibly driven by the pleiotropy of gene networks, repeatedly used in different organs (Liang et al., 2018; Musser & Wagner, 2015).

We further suggest that pleiotropy-induced DSD may explain another observation concerning genes involved in human diseases and pleiotropic genes. It was expected that the embryonic expression profiles of these important genes would evolve slowly, but they evolve as fast as the rest of the genome (Cardoso-Moreira et al., 2019, 2020). Further work may reveal which part of sequence and expression divergence which is usually attributed to genetic drift (divergence by random chance) could in fact be attributed to “pleiotropy-induced DSD”.

## MATERIAL AND METHODS

### Data analysis

R scripts corresponding to the main methods and processed data are available on GitHub (https://github.com/msemon/DriftHamsterMouse).

### Rodent breeding and embryo sampling

CD1 (CD1) adult mice and RjHan:AURA adult hamsters were purchased from Charles River (Italy) and Janvier (France) respectively. Females were mated overnight and the noon after morning detection of a vaginal plug or sperm, respectively, was indicated as ED0.5. Other breeding pairs were kept in a light-dark reversed cycle (12:00 midnight), so that the next day at 16:00 was considered as ED1.0.

The *Bmper^tm1Emdr^* strain (Zakin et al., 2008) was kept in a C57/BL6N background by crossing heterozygotes with wild types, as homozygotes die at birth. To avoid suffering at birth, we generated homozygotes and wild type samples for X-ray by crossing heterozygotes and sacrificing pregnant mice at 19.5 days (1 day before delivery). Pregnant mouse females were killed by cervical dislocation. Hamster females were deeply anesthetized with a ketamine-xylasine mix administered intraperitoneally before being killed with pentobarbital administered intracardially. All embryos were harvested and thereby anesthetized on cooled Hank’s or DMEM advanced medium, weighted as described in (Peterka et al., 2002) and immediately decapitated.

This study was performed in strict accordance with the European guidelines 2010/63/UE and was approved by the Animal Experimentation Ethics Committee CECCAPP (Lyon, France, APAFIS#27308-2020092210045896 v1).

### Estimating embryonic age from embryo weight

Embryo weight is well correlated with developmental age, allowing us to use it as a proxy in mouse and hamster, following (Pantalacci et al., 2009). We fitted age of development according to weight (in mg) for hamster and mouse data separately, based on 1047 mouse embryos and 636 hamster embryos respectively, collected over more than 15 years of research. We fitted generalised additive models (GAM) to the data after Box-Cox transformation of weight (libraries mgv version 1.8-35 for GAM and MASS 7.3-53.1 for Box-Cox). These models were prefered to log transformations and linear models, because they allow to treat the data homogeneously between species, and because the relationship is not perfectly linear between weight and age (Figure S1). These models were then used to predict developmental age, based on weight, for all samples used in this study (RNA-seq analysis, cusp patterning analysis, and *in situ* hybridizations for several genes).

### Epithelium dissociations and in situ hybridizations

Complete or hemi mandibles and maxillae were dissected in Hank’s medium and treated with Dispase (Roche) 10mg/ml in Hepes/KOH 50mM ph7.7; NaCl 150 mM at 37 °C for 30 min to 1h depending on embryonic stage. Epithelium and mesenchyme were carefully separated and fixed overnight in PFA 4% at 4 °C. DIG RNA antisense mouse *Fgf4*, *Shh*, *Fgf10* (Bellusci et al., 1997), *Bmper/Cv2* probes were prepared from plasmids described elsewhere (Coffinier et al., 2002). Mouse *Dkk1*, *Wif1*, hamster *Bmper* probes, Mouse and hamster *Bmp4* probes were newly cloned following RT-PCR or DNA synthesis (Table S1). *In situ* hybridizations were done according to a standard protocol (DIG mix, DIG antibody and BM purple were purchased from ROCHE). Photographs were taken on a Leica M205FA stereomicroscope with a Leica DFC450 digital camera (Wetzlar, Germany) or on a Zeiss LUMAR stereomicroscope with a CCD CoolSNAP camera (PLATIM, Lyon, France).

### Immunolocalisation and 3D reconstructions

Tooth germs dissected from litter-mate embryos of RNA-seq samples were fixed overnight in PFA4% and dehydrated through a methanol series. In toto immunolocalization protocol was adapted from (Ahnfelt-Rønne et al., 2007). Following incubation in H202 5%, DMSO 10% in methanol for 4 hours, they were rehydrated, blocked and incubated successively with a pan-cytokeratin antibody (overnight,1/200, Novus Biologicals) and a Dylight 549 conjugated Donkey Anti-rabbit antibody (overnight 1/200, Jackson immunoresearch) and finally with Hoechst (overnight, 50μg/ml). Following methanol dehydration, they were clarified and mounted in BABB as described in (Yokomizo & Dzierzak, 2010). They were imaged with a Zeiss LSM710 confocal microscope at the PLATIM (Lyon, France). The outline of the epithelium labelled by the pan-cytokeratin antibody and the outline of the tooth germ labelled with hoechst were delineated manually and reconstructed in 3D in the AMIRA software.

### X-ray scanning and 3D reconstruction of *Bmper* and wild type tooth shape at 19.5 days

We obtained 14 homozygote (Ho) and 19 wild type (Wt) samples from a total of 14 different 19.5 dpc litters, out of which we selected for reconstruction of the first molar morphology 7 Ho and 4 Wt with matching body weights (homozygotes: 1174-1329 mg; wt: 1227-1310 mg). At 19.5 dpc, female embryos are more developmentally advanced than male embryos of a similar weight, therefore sex was also recorded. This was necessary to control for differences in growth advancement, since we anticipated (and confirmed) that supplementary cusps are still growing rapidly at this stage, due to their late formation. Heads freed of skin were fixed in PFA, dehydrated in ethanol, stained with 0,3% PTA in 70% ethanol for 2 weeks-1 month and scanned at 40kV on a Phoenix Nanotom S microtomography for a voxel size of 4 µm. Semi-automatic reconstruction of the enamel-dentin junction was performed with ITK-snap. Reconstructions were oriented for comparison in MeshLab 2021.05. Due to variations in staining efficiency and advancement of mineralization, only a total of 4 Ho and 4 Wt upper molars and 3 Ho and 4 Wt lower molars were finally successfully reconstructed and considered to be directly comparable. The aberrant upper molar morphology was obvious on microCT sections in 7/7 samples. The loss of one cusp was observed in 4 Ho/4 3D reconstructions. The larger lingual cusps were observed in all Ho 3D reconstructions when paired with a wt of corresponding age. This (as well as cusp loss) was also confirmed by comparing epithelial dissociations of *Bmper* Ho and Wt embryos at 19.5 dpc.

### Modelling and comparing cusp patterning dynamics

To compare the dynamic of crown morphogenesis in four teeth (lower and upper molars in hamster and mouse) we need to establish the sequence of primary and secondary signalling centres formation (respectively, PEK and SEK). In mouse, this could be done with time lapse imaging of fluorescent lines (Harjunmaa et al., 2014). To integrate non-model species like hamster, we had to set up a new method that infers the dynamic based on fixed embryos. We hybridised developing molars against a *Fgf4* probe to reveal PEK and SEKs. The patterns we observed among samples are consistent with a stereotypic and specific sequence of SEK patterning in each tooth and species (Figure 1B, schemas on the sides). We name each stage by the number of signalling centres (PEK stage then 1-SEK stage, 2-SEK stage etc).

Cusp patterning can be seen as a succession of irreversible stages representing step-wise cusp additions. Transition rates between these stages were modelled through continuous time Markov modelling as in (Pantalacci et al., 2017). The rationale is that if sampling is uniform over the time course of tooth morphogenesis, stages that are rarely sampled are very transient (with high exit rate), while stages that are often sampled last for a longer period of time. In continuous Markov models, the duration of each state follows an exponential distribution, which is not realistic for the stage lengths. So, to have a more realistic stage length distribution, each stage was modelled by several consecutive states, so that its length followed an Erlang distribution, which has a mode different from zero. We built independent models for each species and tooth types. Models are estimated on 121 embryos for the hamster lower molars, 113 for hamster upper, 217 for mouse lower, 187 for mouse upper.

We estimated the duration of each stage in a complete model, with different transition rates for all stages. We also fitted several simpler, nested models, with constraints on the number of different transition rates, up to the most simple model with the same transition rate for all stages. We retained models with three different rates in mouse, and two different rates in hamster, by comparing the fit of the models by likelihood ratio tests in each tooth. Markov models were built by custom scripts calling on R libraries maxLik and expm (maxLik_1.4-8 and expm_0.999-6).

### RNA-seq sample preparation

A total of 32 samples per species, coming from eight individuals, were prepared for the time serie RNA-seq analysis, representing consecutive stages in mouse (ED14.5, 15.0, 15.5, 16.0, 16.5, 17.0, 17.5, 18.0) and nine stages in hamster (ED11.8, 12.0, 12.2, 12.5, 13.0, 13.25, 13.5, 13.75, 14.0). Each sample contained two whole tooth germs, the left and right first molars (M1) of the same female individual, and for a given stage, the upper and lower samples came from the same individual. Harvesting and dissection were performed in a minimal amount of time in advanced DMEM medium. The M1 lower and upper germs were dissected under a stereomicroscope and stored in 200 uL of RNA later (SIGMA). Similarly dissected tooth germs from the same litter and same weight were fixed overnight in PFA 4% for immunolocalization and 3D reconstruction, to check for dissection leaving almost no non-tooth tissue. Examples of dissection are visible in (Pantalacci et al., 2017). Another embryo of the same litter and same weight was processed as indicated above for *Fgf4 in situ* hybridization to check the exact developmental stage. Total RNA was prepared using the RNeasy micro kit from QIAGEN following lysis with a Precellys homogenizer. RNA integrity was controlled on a Bioanalyzer (Agilent Technologies, a RIN of 10 was reached for all samples used in this study). PolyA+ libraries of the large-scale dataset were prepared with the Truseq V2 kit (Illumina, non stranded protocol), starting with 150 ng total RNA and reducing the amplification step to only 12 cycles and sequenced on an Illumina Hi-seq2000 sequencer (100 bp paired end reads) at the GENOSCOPE (Evry, France).

For the bucco-lingual dataset, we dissected the 4 first molars (left/right, lower/upper) from a unique mouse E15.0 embryo (weight: 359 mg) as above, except that tooth germs were cut in two halves to isolate buccal and lingual side. Replicates were thus obtained by comparing the right and left side of this same embryo. Total RNAs were extracted and libraries were prepared as above, starting with 50-70 ng total RNAs, where an equal amount of AmbionR ERCC RNA Spike-In Mix1 had been added according to the AmbionR protocol (e.g. 1μL og a 1 :1000 dilution for each tube). A total of 8 libraries were sequenced (50bp single-end reads) by the Genomeast Sequencing platform, a member of the France Genomique program.

For the epithelium-mesenchyme dataset, lower and upper mouse and hamster first molars were dissected as above and treated for 15 minutes at 37°C with Dispase (Roche) 10mg/ml in Hepes/KOH 50mM ph7.7 ; NaCl 150 mM to separate the epithelial from mesenchymal parts which were stored in RNAlater. For the mouse data, we generated samples for 2 stages in 2 replicates, using embryos from the same litter (stage 15.0 dpc, weight: 350 and 370 mg; stage 16.5 dpc: weight: 788 and 808 mg). Left and right epithelium or mesenchyme were pooled. For the hamster data, we generated samples for a single stage without replication. We pooled the left epithelial or mesenchymal parts from 2 embryos from the same 12.5 dpc litter (413 and 427mg). A total of respectively 16 and 4 libraries were generated with Truseq V2 kit and sequenced (50bp single-end reads) by the Genomeast Sequencing platform.

### Multivariate analyses

Multivariate analyses were performed using the ade4 package (ade4_1.7-18; (Dray & Dufour, 2007)). We performed principal component analyses on normalised counts (DESeq basemeans), and between groups analyses on the resulting components, which allowed us to quantify the proportion of variance associated with each factor.

We used RAPToR (RAPToR_1.1.4, (Bulteau & Francesconi, 2021)) to estimate the offset between upper and lower molar development. We staged upper molar samples on reference made from lower molar samples conjointly for both species. Interpolations were done with gam models fit on 5 components of an ICA.

### Expression levels estimation using RNA-seq and differential expression analysis

For the whole tooth germ data (64 samples) we obtained 100 bp paired-end sequences, with on average 46.2M (millions) reads per sample. For epithelium/mesenchyme and bucco/lingual data (respectively 20 and 8 samples), we obtained 50 bp single-end sequences, with on average 93.7M and 48.6M reads per sample respectively. Raw data are publically available in ENA with project accession number: PRJEB52633.

These reads were mapped by using Kallisto (version 0.44.0, (Bray et al., 2016)) to custom reference sequences for hamster and mouse transcriptomes. To generate them, we retrieved mouse and hamster cDNAs from Ensembl (release 93, July 2018, assemblies GRCm38.p6 and MesAur1.0 (Howe et al., 2021)), selected 14536 pairs of one-to-one orthologous transcripts, realigned pairs of sequences with Macse (macse_v2.01, (Ranwez et al., 2011)) and cropped the alignments to get orthologous segments by using custom scripts to make expression levels comparable between species.

Differential gene expression analysis (DE analysis) was performed on smoothed expression profiles over developmental age. Developmental age was estimated with embryo weight (GAM models above) and stages were homologized based on morphological criteria at early cap stages and late bell stage (14.5/18.0 days in mouse; 12.2/14.5 days in hamster). These boundaries, confirmed by PCA analysis of the transcriptome data (Figures 1 and S1) were used to convert days of development into relative development age (0-10).

Expression profiles were fitted by third degree polynomial splines with 2 interior knots, for each tooth and species (bs function of spline R package (W. Wang & Yan, 2021), independently or jointly within tooth and/or species, as explained below. Nested models were tested by DEseq2 (Love et al., 2014) and the best model was chosen for each gene by comparing the fit of these nested models (FDR adjusted p-value < 0.05 from DESeq2 LRT tests). When we compared temporal profiles between species, we accounted for the average level of expression in each species. This is to focus on changes in regulation over development, and to discard potential remaining artefacts in species-specific quantifications. Several tests were performed and are described below with the corresponding figure number.

To model temporal expression profiles in each species separately (Related to Figure 2A), we compared a “simple” model with a single curve for both time series to a “complex” model with one specific curve per tooth. This procedure does not directly provide the sense of the bias. To estimate it in each species, we computed the values predicted by the upper and lower models for 100 equally distributed points (i) on the timeline, and measured the distance point by point as follows : sum((up(i)-low(i))/(up(i)+low(i)) (Mus.dist.sign or Ham.dist.sign in Table S2). This measure evaluates whether overall, expression levels in upper molar are above or below lower molar’s.

To model the divergence of temporal expression profile in each tooth type separately (Related to Figure 6D), we compared a “non-divergent” model with a single curve to fit both time series (with a species-specific offset to only consider the temporal dynamic), to a “divergent” model with one specific curve per species (with a species-specific offset).

To model temporal expression profiles in the 4 tooth types with tooth-specificity in one species (related to Figure 3B), the four time series were modelled as in the “hamster/mouse”, plus “mouse tooth-specific” and “hamster tooth-specific” models, which correspond to distinct curves for upper and lower molars in one species, but not in the other. Genes were selected by comparison with the “hamster/mouse” model above.

Selection of the temporal expression profile in the 4 tooth types was done as follows (related to Figure 6A). The “simple” model fits a single curve for the four time series. The “complex” model fits 4 different curves, one per tooth type. The “hamster/mouse” model has 2 different curves, one per species. The “upper/lower” model has one curve per tooth, including the species-specific offset. The best model was selected for each gene by using a bottom-up approach with the results of four independent tests: t1 compares “hamster/mouse”*versus* simple model; t2: “upper/lower” *versus* simple; t3: complex *vs* “upper/lower”; t4: complex *vs* “hamster/mouse”. If t1 and t2 are not significant, then the simple model is chosen. If t1 is significant and not t2, the gene is assigned to: “hamster/mouse”. If t2 but not t1: “upper/lower”. Finally, if “lower/upper” or “hamster/mouse” and t4: complex.

From this selection procedure, percentage of coevolution among genes was computed as the proportion of selected “hamster/mouse” models among the selected models as follows (related to Figure 6A): “hamster/mouse”/(“hamster/mouse”+“upper/lower”+“complex”).

We then computed the intersection of the results with several lists of genes important for tooth development: 259 genes from the bite-it database (retrieved in July 2019), 187 genes with a phenotype in tooth development (100 “dispensable” genes, 87 “keystone” genes, (Hallikas et al., 2021)), and 295 genes belonging to 8 pathways active in tooth development (17 genes in ACTIVIN pathway, 81 in BMP, 10 in EDA, 69 in FGF, 32 in SHH, 9 in NOTCH, 11 in TGFB, 96 in WNT, courtesy Jukka Jernvall).

### Gene Ontology analysis

Gene ontology (GO) analysis was performed and visualised with gprofiler2 (gprofiler2_0.2.0, (Kolberg et al., 2020)), clusterProfiler (clusterProfiler_3.18.1, (Wu et al., 2021)), and simplifyGO (simplifyEnrichment_1.2.0 (Gu & Hübschmann, 2021)) using the full list of genes expressed in the corresponding dataset as a background.

### Measure of pathway activation in RNAseq samples

ROMA was used to quantify activation of WNT, BMP and SHH pathways in the bucco-lingual samples (version rRoma_0.0.4.2000, https://github.com/Albluca/rRoma and (Martignetti et al., 2016)). ROMA is designed to compare pathway activity in transcriptomic samples based on expression levels of a list of targets for the pathway. Genes for the SHH modules were retrieved from GSEA ((Mootha et al., 2003), 41 genes present in our dataset). Because BMP and WNT pathways are active both in the mesenchyme and the epithelium and they target different genes in each tissue (O’Connell et al., 2012), we used two separate lists of targets to estimate both an epithelial and a mesenchymal activity, adapted from a “regulatory evidence” dataset established for first lower molar development (O’Connell et al., 2012). Building on literature and their own transcriptomic analysis, the authors had defined target genes based on their up or downregulation following activation or inactivation of each pathway. For data consistency, we selected only targets established in the study from transcriptome analysis, in 13.5 and 14.5 dpc epithelium and 10.5 dpc mesenchyme. Different modules were built for activities in the mesenchyme and epithelium compartments. For BMP in the mesenchyme, we considered 15 genes as positive targets and 4 as negative targets (further noted 15:4). In the epithelium, the numbers of positive:negative targets were respectively 32:34. For WNT, we built modules with 4:31 positive:negative targets in the mesenchyme, and 33:45 in the epithelium. These in-silico quantifications were consistent with many known aspects of tooth development. Buccal compartments all show high levels of pathway activity, consistent with the presence of the first SEK acting as a source of WNT, BMP and SHH signals. Lingual compartments show much lower levels of signalling activities than buccal compartments, consistent with their distance to the first SEK. The lower lingual compartments show BMP and WNT activities that are higher in epithelium than in mesenchyme, consistent with the fact that epithelial activation predates mesenchymal activation in tooth development.

### Estimating tissue proportions from RNA-seq data by deconvolutions

We used the R package DeconRNASeq (DeconRNASeq_1.32.0 (Gong & Szustakowski, 2013)) to estimate the relative proportions of epithelium and mesenchyme compartments in bulk tooth germ transcriptomes. We defined lists of marker genes for each compartment by pairwise differential analysis of tissue-specific transcriptomes (DESeq2, log2 fold change > 3, adjusted p-value < 0.05). We used 1025 mesenchyme and 621 epithelium marker genes found by comparing 10 epithelium and 10 mesenchyme RNAseq samples, mixing tooth, stages and species. We estimated the accuracy of the prediction by bootstrapping 1000 times the marker lists. The relative proportions of buccal and lingual compartments was inferred by a similar procedure. We used 414 buccal and 235 lingual marker genes, from the differential analysis of 8 samples (DESeq2, log2 fold change > 1, adjusted p-value < 0.05).

### Expression levels and transcriptome dynamics in bats

We downloaded all bats and mouse raw RNA-seq samples from a published dataset (SRP061644, Maier et al. 2017), totalizing 17 samples in mouse and 16 in bat (*Carollia Perspicillata*) at three consecutive stages: ridge (CS13), bud (CS14) and paddle (CS15) stage. Bat reads were assembled *de novo* with Trinity v2.14.0 (Grabherr et al., 2011), by using single end mode and *in silico* normalisation. Bat expression levels were quantified by Salmon (Patro et al., 2017) with the script provided by Trinity (align_and_estimate_abundance.pl). Mouse reads were directly mapped with Salmon to the GENCODE mouse transcriptome reference (gencode.vM29.pc, (Frankish et al., 2021)). Bat transcripts were assigned to mouse orthologs by blastn (Altschul et al., 1990). Blast and Trinity were run with prebuilt dockers. Differential analysis was run over smoothed expression profiles like in the method section “Expression levels estimation using RNA-seq and differential expression analysis”. Code is available here: https://github.com/msemon/DriftHamsterMouse

## ACKNOWLEDGEMENTS

We acknowledge the contribution of several platforms of SFR Biosciences Gerland-Lyon Sud (UMS344/US8): the Plateau de Biologie Expérimentale de la Souris (PBES) (many thanks especially to Jean-Louis Thoumas, Tiphaine Dorel, Céline Angleraux, Marie Teixeira, Myriam Prudent), the Plateau Technique Imagerie/Microscopie (PLATIM) and the ANIRA/IMMOS platform with Mathilde Bouchet; as well as the computer resources from PSMN (ENS Lyon). We acknowledge the technical help of Anne Lambert, Alain Rubod, Mathilde Estevez-Villar, Claudine Corneloup and the contribution of many students including Coraline Petit, Alice Lorenc, Margaux Pillon, Ludivine Rotard and Asma Benahmed. We are grateful to several colleagues and their staff for sending plasmid probes: Irma Thesleff (*Fgf4)*, Hiko Ogura (*Cv2/Bmper)*, D. Duboule (*Fgf10*). We are grateful to Pamela Lockyer, Cam Patterson and Xinchun Pi for collecting our very first *Bmper* embryos. We are grateful to Jennifer Esser for kindly providing the *Bmper* strain and advice on genotyping. We kindly thank Mirko Francesconi, Marie Delattre, Michaelis Averof, Pascal Hagolani for their feedback on the manuscript.

This work was supported by the Agence Nationale pour la Recherche (ANR 2011 JSV6 00501 “Convergdent”), a grant from the GENOSCOPE - Centre National de Séquençage and a grant from IDEX Lyon ANR-16-IDEX-0005. Salaries were supported by the Centre National de la Recherche Scientifique, the Ecole Normale Supérieure de Lyon and the Université de Lyon, Université Lyon 1. Klara Steklikova benefited from a Barrand Fellowship program.

