## Supplemental Material (pdf) for "Phenotypic innovation in one tooth induced concerted developmental evolution in another"

### SUPPORTING INFORMATION

#### List of supporting tables

S1 Table: primers and sequences used to synthetize probes

S2 Table: an excel file with results of the tests performed in figure 2A and 6D.

#### List of supporting figures from Fig S1 to S12

Figure S1: Comparing dynamics of morphogenesis and gene expression across teeth and species.

Figure S2: Temporal profiles of a selection of transcription factors with differential expression in both mouse and hamster molars

Figure S3: Simplified GO enrichment results of the genes with a different profile in upper versus lower molar and a similar bias in hamster and mouse.

Figure S4: Transcriptomes series show a developmental delay of the upper molar in both hamster and mouse.

Figure S5: Dynamics and pattern of lingual and posterior cusp addition during early development of lower and upper molars in mouse and hamster.

Figure S6: Simplified Gene Ontology (GO) enrichment for the lists of genes with a specific profile in hamster upper/lower molars (A) or in mouse upper/lower molars (B).

Figure S7: *Sfrp2* and *Osr2* transcriptomic profiles

Figure S8: Bmper spatio-temporal pattern from in situ hybridizations.

Figure S9: The expression changes marking the initiation of cusp patterning are anticipated in mouse molar mesenchyme.

Figure S10: Mouse molars show a precocious transition to SEK stage, with more focalized expression of signalling molecules.

Figure S11: *Shh*, *Fgf8* and *Grem1* co-evolved in the mouse/bat transcriptome dataset

Figure S12: Transcriptional profiles for *Fgf3* and *Inhba* (coding for ACTIVINβA) suggest that they are not involved in the mouse upper molar evolution.

S1 Table

| Probe | size | Primers for RT-PCR/synthetized sequence |
| --- | --- | --- |
| musWif1 | 641bp | Fwd ATCCTACCTTGCCTGCTCCT<br>Rev<br>CAGAAGCCAGGAGTGACACA |
| musDkk1 | 410bp | Fwd TGGCCGTGTTTACAATGATG<br>Rev AAAATGGCTGTGGTCAGAGG |
| mesocricetusBMP4 | 585bp | CTGGTAACCGAATGCTGATGGTCGTTTTATTATGCCAAGTCCTGCTAGG<br>AGGCGCGAGCC<br>ATGCTAGTTTGATACCTGAGACCGGGAAGAAAAAAGTCGCCGAGATTC<br>AGGGCCACGCGG<br>GAGGACGCCGCTCAGGGCAGAGCCATGAGCTCCTGCGGGATTTTGAG<br>GCGACACTTCTGC<br>AGATGTTTGGGCTGCGCCGCCGTCCGCAGCCAAGCAAGAGCGCCGTC<br>ATTCCGGATTACA<br>TGAGGGATCTTTACCGGCTCCAGTCTGGGGAGGAGGAAGAGGAAGA<br>GCAGAGCCAGGGAA<br>TGGGGCTGGAGTACCCCGAGCGTCCAGCCAGCCGGGCCAACACTGT<br>GAGGAGTTTCCATC<br>ACGAAGAACATCTGGAGAACATCCCAGGGACCAGTGAGAACTCCGCC<br>TTTCGTTTCCTTT<br>TCAACCTCAGCAGCATCCCAGAGAATGAGGTGATCTCCTCTGCGGAG<br>CTCCGCCTGTTTC<br>GGGAGCAGGTGGACCAGGGCCCCGACTGGGAGCGGGGCTTCCACC<br>GGATCAACATTTATG<br>AAGTTATGAAGCCCCCAGCAGAAATGGTGCCTCGGCACCTCATCA |
| musBMP4 | 585bp | Fwd: CTGGTAACCGAATGCTGATGGTC<br>Rev:TGATGAGGTGTCCAGGAACCATT |
| mesocricetusBmper | 601 | GTCCTTCATTGGCTCTTCCATGTAAGTGTCAGAGACACTAGATTTCAAA<br>GTCTATGGACTAGAACAAAGGTCATCGCTGAGGACAGAGGACTGGCT<br>TGATGCACCTCCCCTTATGAAGAACCAGATTTGCTGGGCAGTGGCAGC<br>CTGCAATACATGCCTTATTACACGGGCCAATCTCGTTCCAGTTGTCACA<br>GGTCTTTACACATCCAGGGCCACAGGTATCATAAACAGCGCCATGTTT |

| | | |
|---|---|---|
| | | GCACTGGGTAGCTGCACAGCTCTGCTGAGGCTCCCAGTGCACTTTGAT<br>GCCCTCTCTCTGGCAGGCACGGGTATATGCCAGAAACGACTCACAATA<br>ACAGTTTTTATGGACTGGACATTCGCACATGTCTGTCACACAGGACCG<br>GTAGAAAGTGGTATAGTCCACCGTTGAGTGGCAGGTCTGGAACTCCC<br>AGGACTTGAGTTTCTGGCATTCTCGGTGGGCTCGAAGCTTCACCTTCA<br>CTGTGCCTTGACATAGCTCAGGCACTGGTTTTCTCTGTGGTCTGTTGCA<br>GAACTCATTGGACTCCACCCTCCATGACTCAGCAAAGTCATCCACATC<br>AAACTTGAAGTTCCCATCTCCACC |

S2 Table: an excel file with results of the tests performed in figure 2A and 6D.

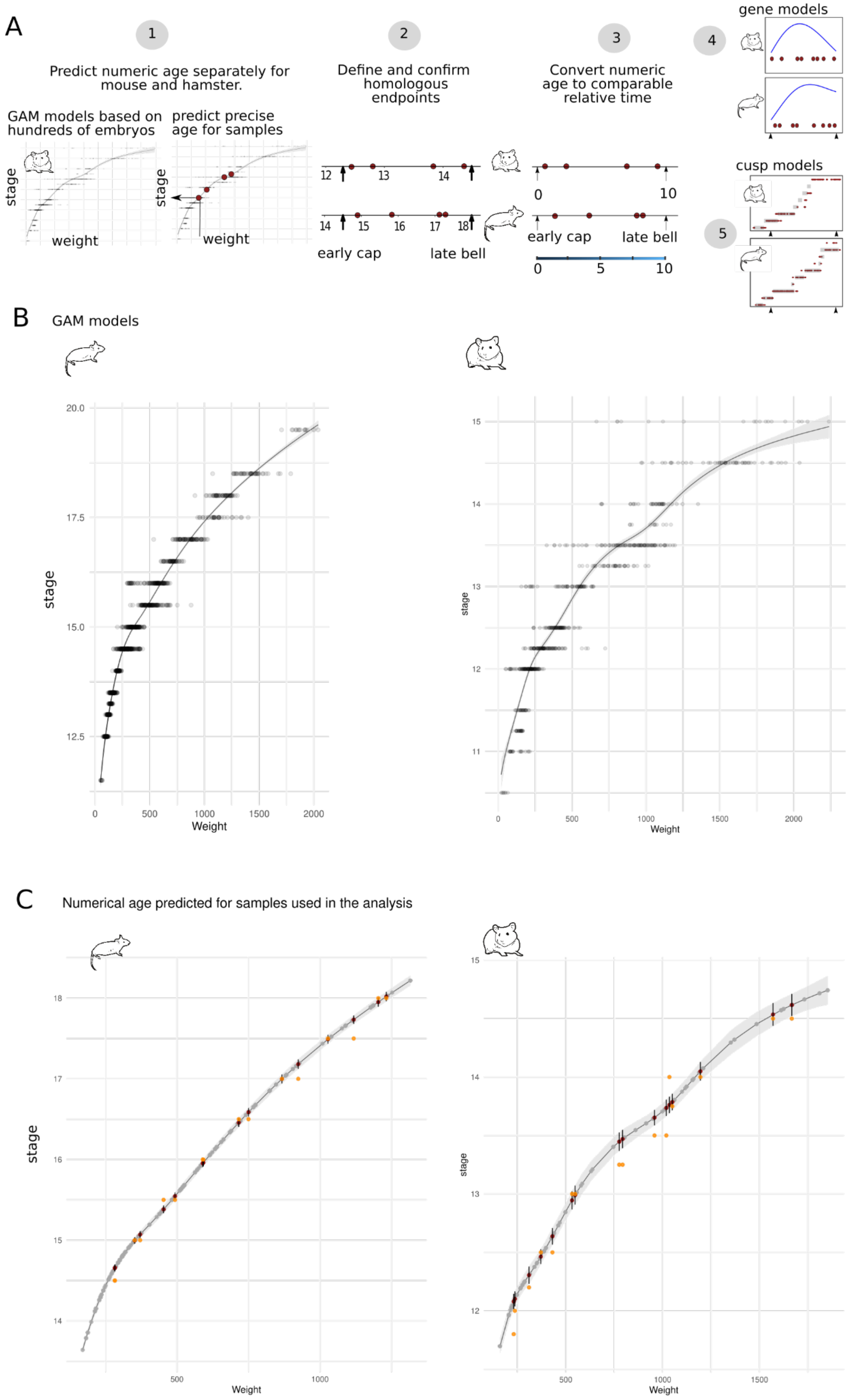
A
1
Predict numeric age separately for mouse and hamster.
GAM models based on hundreds of embryos
predict precise age for samples
stage
weight
2
Define and confirm homologous endpoints
12 13 14
14 15 16 17 18
early cap
late bell
3
Convert numeric age to comparable relative time
0 10
early cap
late bell
0 5 10
4
gene models
5
cusp models
B
GAM models
stage
Weight
C
Numerical age predicted for samples used in the analysis
stage
Weight

**Figure S1 | Comparing dynamics of morphogenesis and gene expression across teeth and species.**

To compare the gene expression dynamics and cusp patterning we wish to work on the fine-scale dynamics rather than comparing samples taken at supposedly homologous stages.

A: We estimate embryos’ developmental age from their body weight through a model established by using hundreds of embryos in each species. Numeric age is predicted separately for mouse and hamster (1). Homologous start and end points are defined based on morphology and confirmed by RNA-seq (Figure 1C) (2). Numeric ages are rescaled to comparable relative time of development, from 0 to 10 in each species (3). Relative times are estimated for the 64 RNA-seq samples used to build gene temporal profiles using polynomials (4). Relative times are also estimated for a series of fixed embryos that were hybridised against a *Fgf4* probe to reveal PEK and SEKs. The relative duration of each stage was modelled with continuous Markov processes which assume that very few embryos should be sampled at very transient stages, while many at longer stages.

B: Relationship between embryonic stage (days post coitum, dpc) and weight (mg) was fitted in each species by a GAM model on boxCox transformed values, based on 1047 mouse and 636 hamster embryos respectively.

C: Predicted developmental ages are shown for the samples used to model cusp patterning (grey points) and for the samples used in RNA-seq analysis (whole tooth germs, dark red points). Confidence intervals (95% percentiles) are indicated for each prediction. Days post coitum are indicated in orange for samples used in RNA-seq analysis, for comparison.

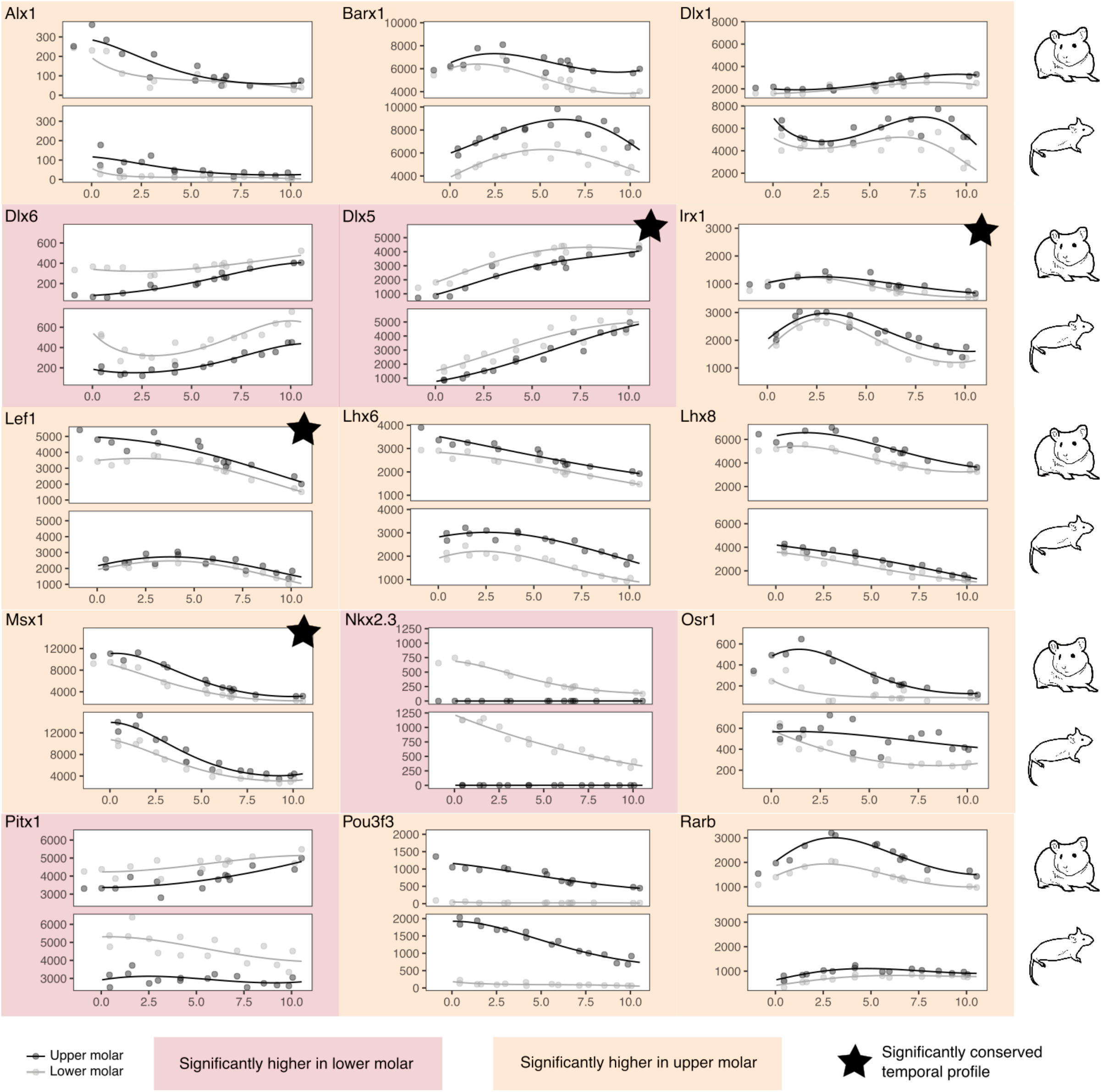


**Figure S2: Temporal profiles of a selection of transcription factors with differential expression in both mouse and hamster molars** (out of a total of 550 genes). Colours highlight genes which expression is consistently higher in lower molars (red) or upper molars (orange) of both species. The black star indicates genes whose temporal profile is conserved between mouse and hamster in both teeth (out of 165 genes).

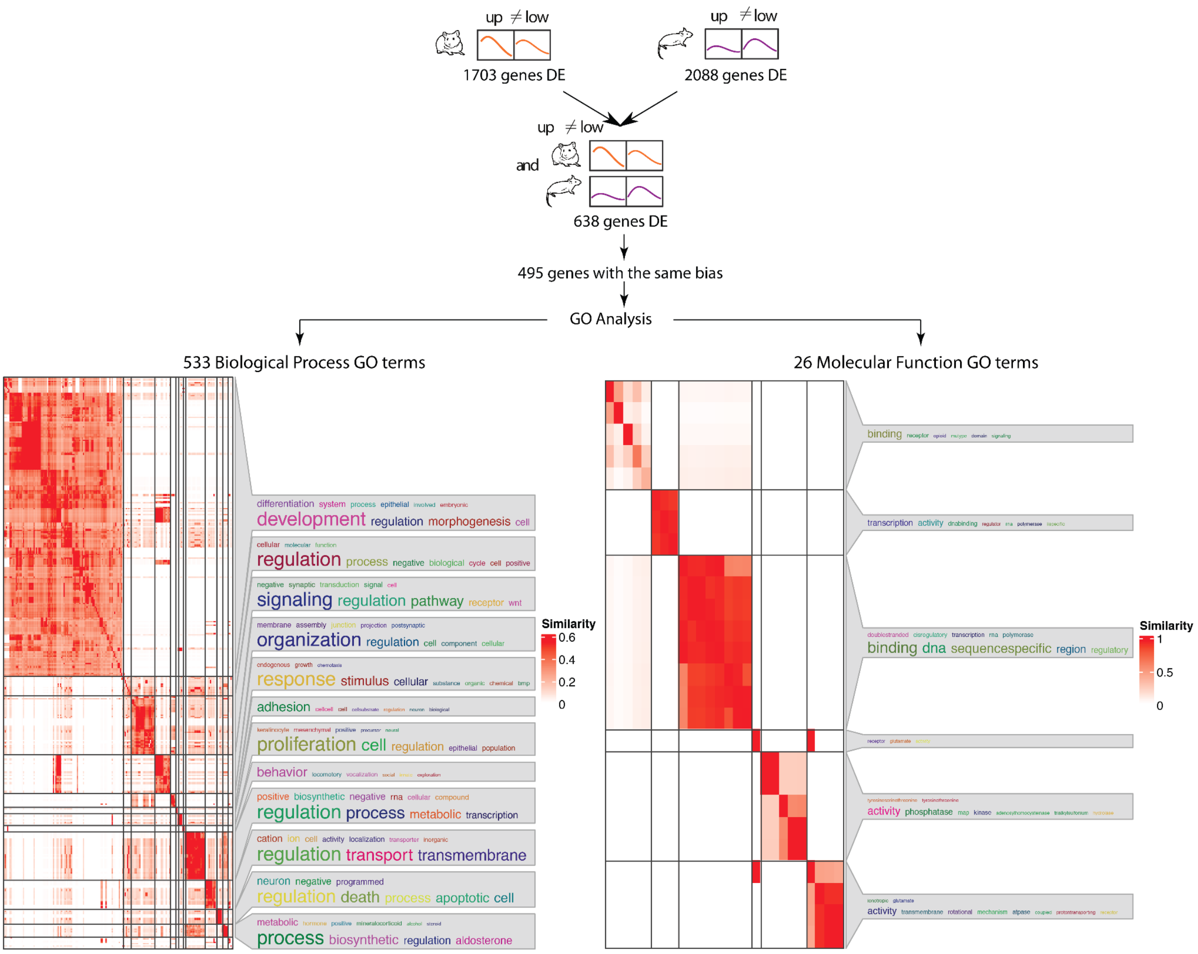


**Figure S3: Simplified GO enrichment results of the genes with a different profile in upper versus lower molar and a similar bias in hamster and mouse.** Genes were detected as in Figure 6. The matrix was drawn by clustering the corresponding semantic similarity matrix of the significant GO terms (simplifyEnrichment, biological process and molecular function gene ontology). On the right side of the heatmap there are the word cloud annotations which summarise the functions with keywords in every GO cluster. Note there is no word cloud for small clusters (size < 5).

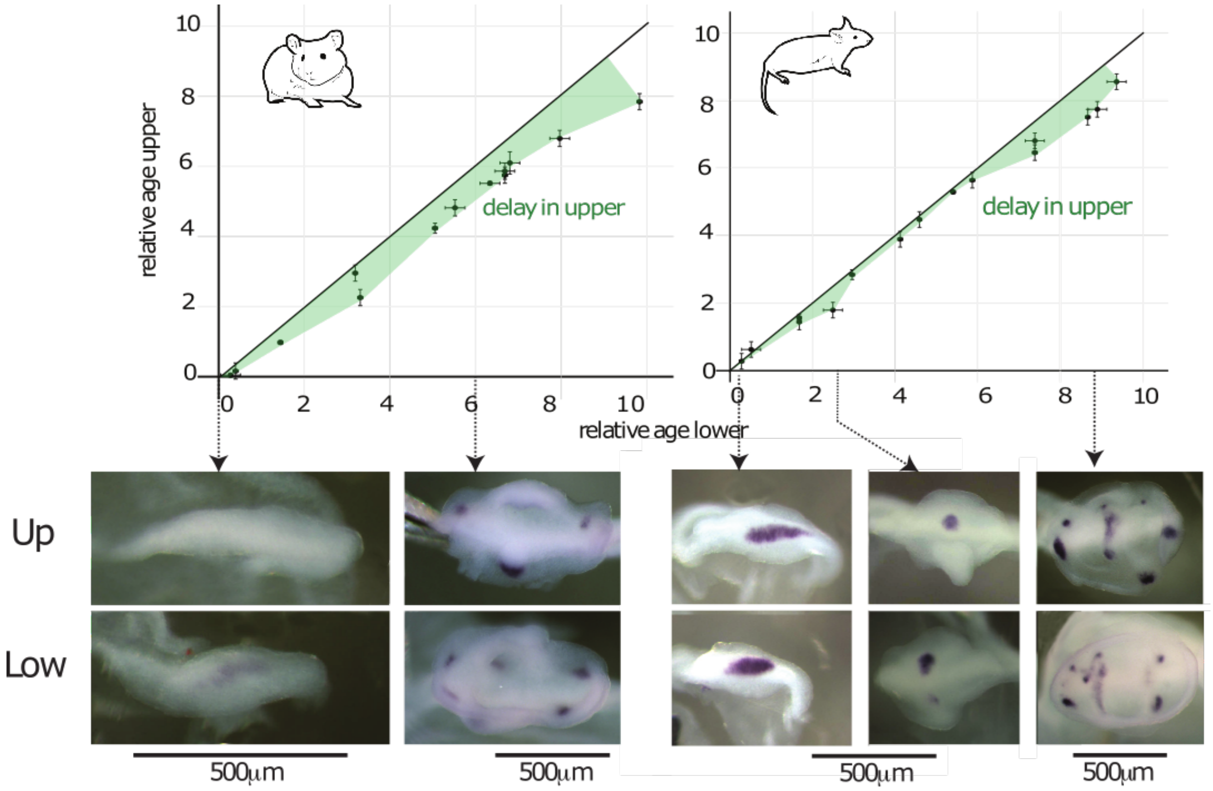


**Figure S4: Transcriptomes series show a developmental delay of the upper molar in both hamster and mouse.**

Relative age of development was measured by RAPToR for upper and lower molars of the same embryos. Upper molars show a systematic delay as compared to lower molars, shown in green. Confidence intervals were obtained by bootstrapping on random subsets of genes. Below, pictures show *Fgf4 in situ* hybridization for the epithelium of lower and upper developing molars taken from the same embryo (age is indicated by arrows and stages in white). Note the obvious delay of upper molar development in hamster: In the youngest embryo, cap transition has occurred in lower molar (PEK stage) but not yet in upper molar (Bud). In the oldest embryo, progression of crown formation (cap shape of epithelium) and cusp formation (6 SEK stained with *Fgf4* versus 4) is obviously more advanced in the lower molar. The situation is less obvious in mouse in the early cap stage (note however that the PEK is already more roundish in the lower molar) but visible when the upper molar has only 1 SEK compared to 2 SEK in the lower molar, or later when cusp patterning is still proceeding in the upper molar but completed in the lower molar.

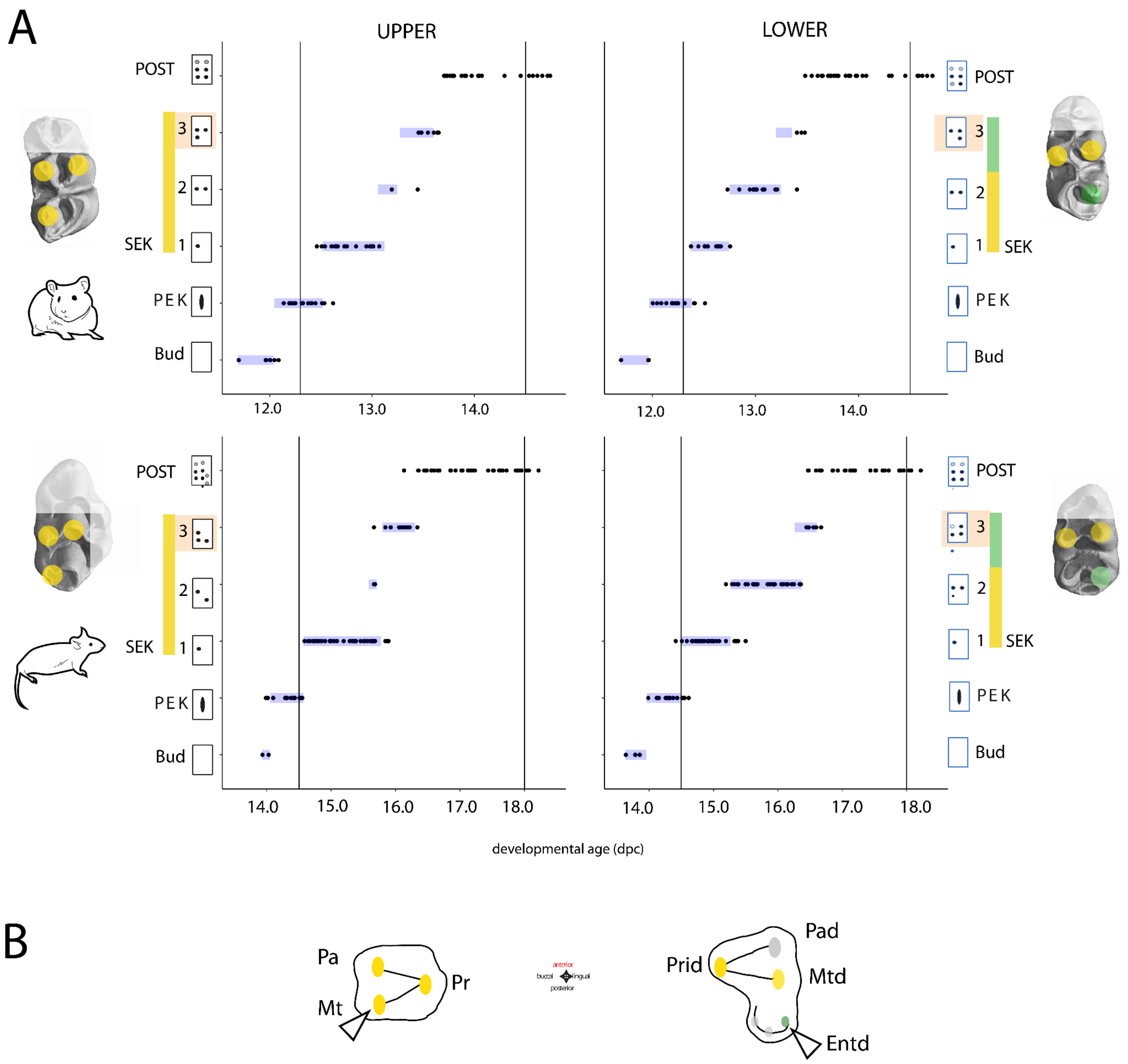


**Figure S5: Dynamics and pattern of lingual and posterior cusp addition during early development of lower and upper molars in mouse and hamster.**

A. To focus on conserved differences in early bucco-lingual/posterior development of lower versus upper molar we modelled the patterning of the same four posterior cusps, discarding information about anterior cusps and supplementary lingual cusps. Each panel represents a series of *Fgf4*-hybridised samples (black dots) with their developmental age and signalling centre stage. Stage duration was modelled using markov processes with three different rates (slow, medium and rapid corresponding to long, medium and short stages). Stage durations are represented in blue, and centred on for each stage on developmental time with maximum probability. Developmental age was estimated through a relationship between embryonic weight and age post coitum (dpc).

B. Protoconid-Prid, Metaconid-Mtd, Entoconid -Entd, Paraconid-Pad and Paracone-Pa, Protocone-Pr, Metacone-Mt, respectively. Stage 3-SEK differs in the two molars, in terms of duration and cusp pattern (longer with buccal cusp in upper molar; shorter with lingual cusp in lower molar). Homology with cusps of the early mammals' tribosphenic molars is shown. The tribosphenic upper molar had 3 main cusps, homologous to the three first-developing cusps in mouse and hamster (yellow). The tribosphenic lower molar had three main cusps (one of which, the paraconid, was lost in basal rodents. The present anterior cusps of mouse and hamster are not homologous to this cusp, as they newly evolved in the lineage formed by mouse and hamster) and a posterior "bassin" formed by shorter cusps and deported lingually to occlude with the lingual protocone of the upper molar. The 3-SEK in mouse and hamster is the entoconid, homologous to the entoconid of the posterior bassin and lingual as him (green dots). It is remarkable that the formation of cusps corresponding to the lower molar tribosphenic bassin is occuring after a long 2 SEK stage in both species, during which the posterior part of the lower molar rapidly grows. This reflects the antero-posterior organisation of the lower tribosphenic tooth with tall main cusps corresponding to pre-mammalian cusps, and a phylogenetically more recent bassin of shallow cusps. The 2 SEK stage is short in upper molars, consistent with the third cusp being one of the three main cusps. In contrast, the 2 SEK stage of the upper molar is short in both species, the three first cusp corresponding to the three main cusp found both in the mammalian tribosphenic tooth and in the pre-mammalian molars.

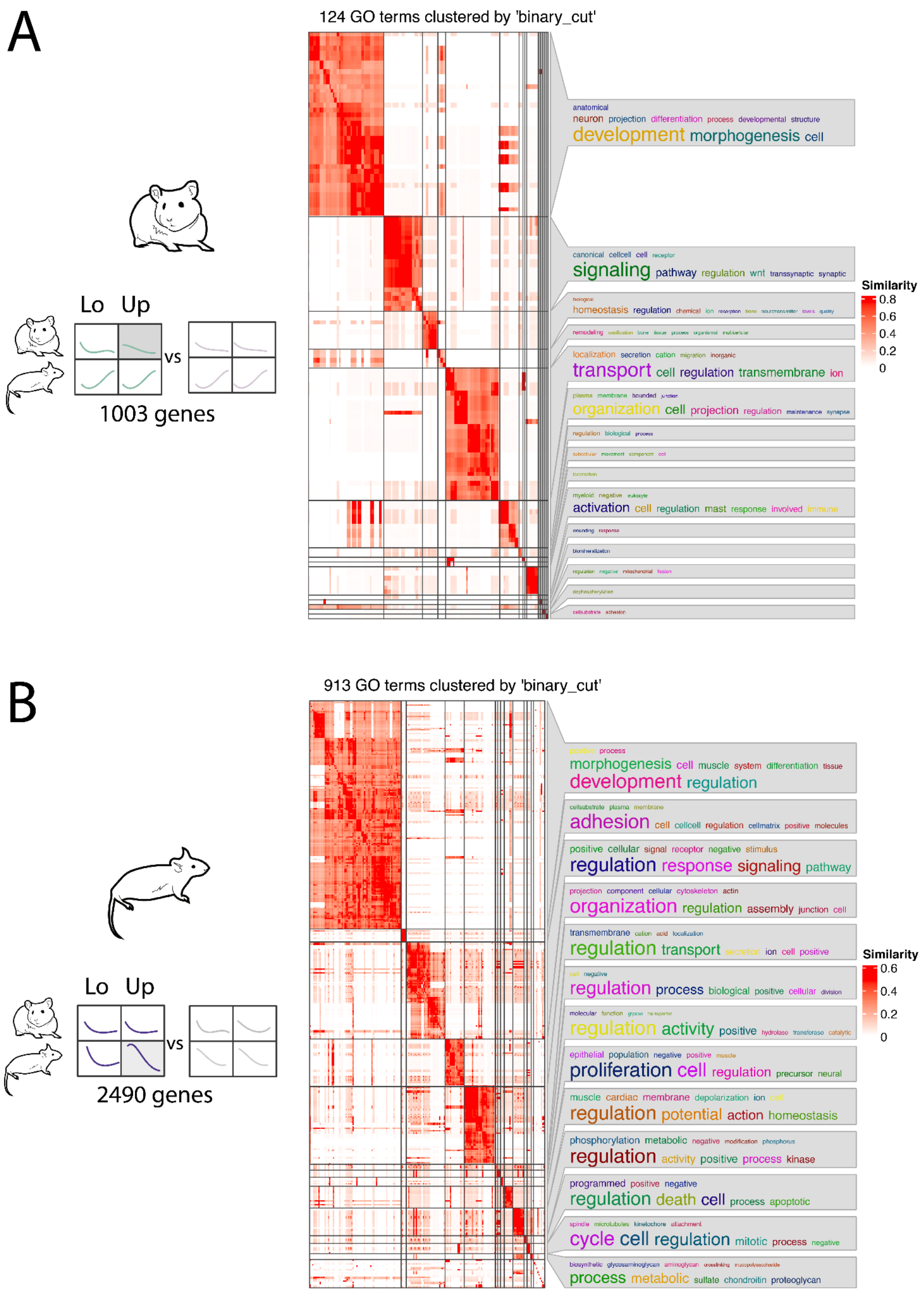


**Figure S6: Simplified Gene Ontology (GO) enrichment for the lists of genes with a specific profile in hamster upper/lower molars (A) or in mouse upper/lower molars (B).** Genes detected as in Figure 2C. The matrix was drawn by clustering the corresponding semantic similarity matrix of the significant GO terms (simplifyEnrichment, biological process Gene Ontology). On the right side of the heatmap, word cloud annotations summarise the functions in every GO cluster with keywords. Note there is no word cloud for small clusters (size < 5 GO terms).

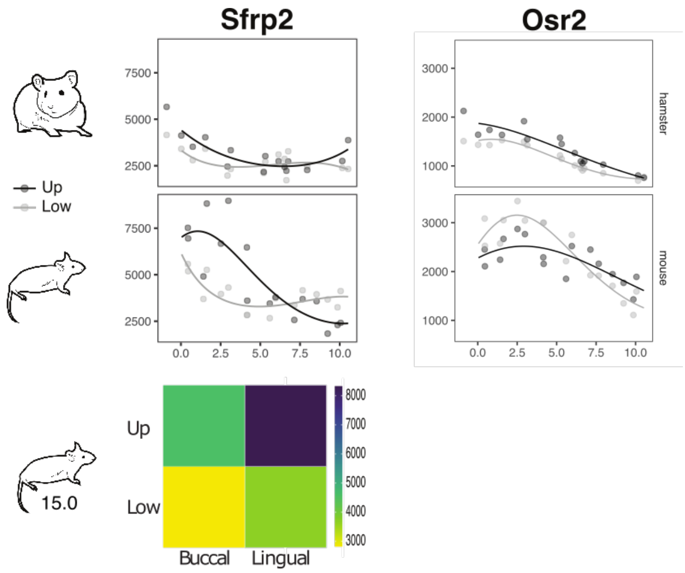


**Figure S7: *Sfrp2* and *Osr2* transcriptomic profiles**

Top: transcriptomic profiles in mouse and hamster. Bottom: spatial profile in the bucco-lingual dataset at 15.0 dpc in mouse. The colour scale represents normalised read numbers for each gene. *Sfrp2* are lingual.

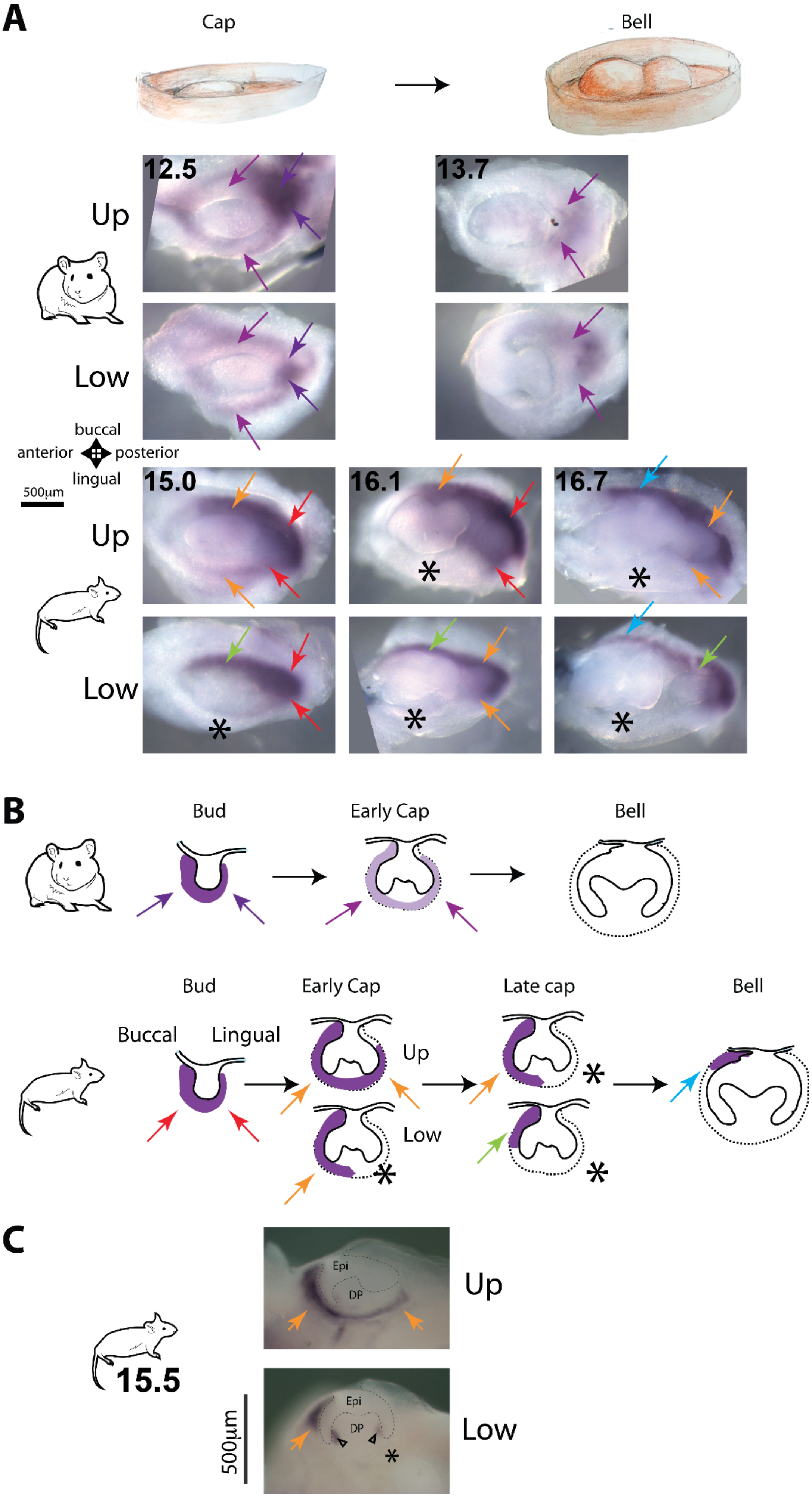
A
Cap
Bell
12.5
13.7
Up
Low
buccal
anterior
posterior
lingual
500µm
15.0
16.1
16.7
Up
Low
B
Bud
Early Cap
Bell
Bud
Early Cap
Late cap
Bell
Buccal
Lingual
Up
Low
C
Epi
DP
Up
15.5
Epi
DP
Low
500µm

**Figure S8: Bmper spatio-temporal pattern from in situ hybridizations.**

A and B- Panel A shows in situ hybridization of jaw mesenchyme with mouse or hamster *Bmper* probes. We studied *Bmper* expression in the mesenchyme of the whole molar row, including the region where the second molar forms, from cap to bell stage of first molar development (note that this region is not included in our transcriptomes, made from dissected first molars only). Because second molar development is delayed, this enabled us to deduce *Bmper* expression from bud to bell stage. Panel B shows schematized sections of developing molars at bud, cap and bell stage with *Bmper* expression deduced from A and C. Colored arrows point to corresponding areas between the jaw mesenchyme (first or second developing molar) and the schematized sections. Note that the dental papilla appears slightly colored by transparency when signal is present around it. Stars point to the absence of signal in the lingual part of the tooth germ. Days of development are indicated in corners. *Bmper* shows a complex spatio-temporal pattern in the tooth mesenchyme, with marked differences between species and teeth. In both species, we found that *Bmper* is strongly expressed in the mesenchyme at bud stage (violet/red arrows), but excluded from the dental papilla which forms with cap stage (purple/orange arrows). In hamster, expression around the M1 cap and its papilla progressively diminishes and has totally vanished in the M1 bell stage. In contrast with this homogeneous decrease, expression in mouse progressively withdraws from the lingual part soon after cap stage (star), but remains high in the buccal part. In the upper molar however, this withdrawal is delayed, and *Bmper* is expressed in a gradient along the bucco-lingual axis.

C Section of 15.5 mouse jaws hybridized with *Bmper*. Colored arrows point signal and star points absence of signal as in cartoons on panel a. Arrowheads point expression around cervical loops exclusively in the very central part of the tooth.

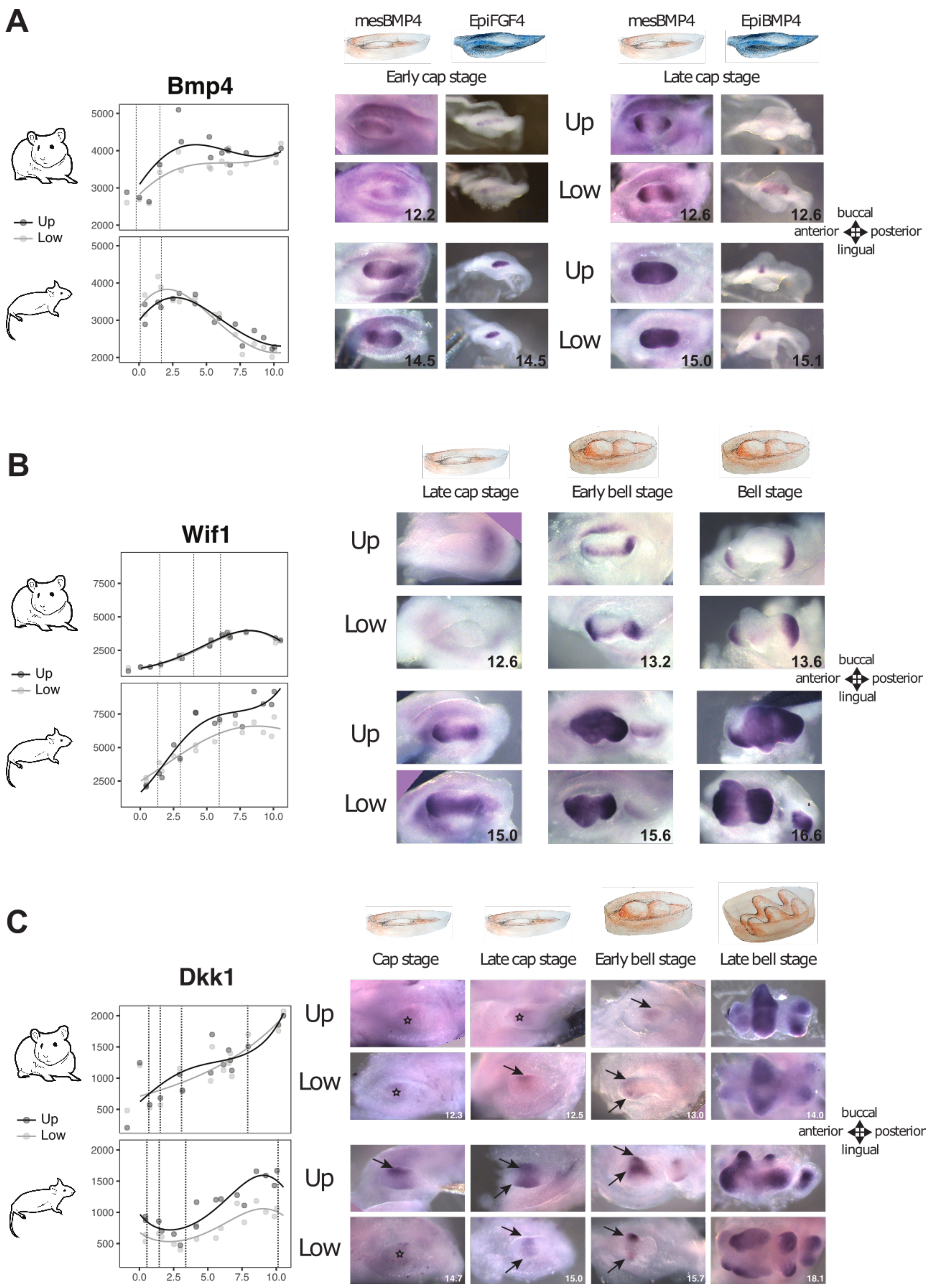


Figure S9: The expression changes marking the initiation of cusp patterning are anticipated in mouse molar mesenchyme.

A,B,C: Left: Transcriptomic profiles for *Bmp4, Wif1* and *Dkk1*. Although *Bmp4* and *Wif1* are also expressed in the epithelium (Figure S9), the transcriptomic profile is mainly driven by the mesenchymal expression representing a larger number of cells. Dashed lines on the profile indicate timepoints of samples shown on the right. Right: top view of the mesenchyme and epithelium, hybridized with the indicated probe. Pictures are centred on the dental mesenchyme, whose initial ovoid shape progresses to a tooth shape through cusp formation.
A: Precocious expression of *Bmp4* in mouse molars. Left: *Bmp4* expression is higher and peaks earlier in both mouse molars. Right: in situ hybridization for a *Bmp4* or *Fgf4* probe, on mesenchymal (left columns) or epithelial (right columns) parts of cap stage tooth germs taken early or late, at similar advancement of epithelial caping for mouse and hamster (compare epithelia). Shortly after cap transition, *Bmp4* mesenchymal expression is low in the hamster, but already strong in the anterior and posterior part of the mouse dental mesenchyme. Later expression is increased in the hamster and resembles early cap stage in the mouse, but in the mouse, *Bmp4* is now seen in the whole dental mesenchyme.
B-Left: *Wif1* expression is higher in mouse and increases earlier as compared to hamster. Right: In situ hybridization of confirmed the more precocious and ubiquitous expression of *Wif1* in mouse.
C: Focused expression of the Wnt inhibitor *Dkk1* is observed earlier in mouse molars, especially in mouse upper molar. Left: *Dkk1* is more highly expressed in mouse upper molars (cusp model as in Figure 4D, adjusted p-value = 0.01). Right: Consistent with the transcriptomic profile, *Dkk1* expression is higher in the upper mouse molar relative to its lower counterpart throughout development. At early cap stage, *Dkk1* expression is well focused (arrow) buccally in the mouse upper molar, where the first upper molar cusp forms. It is more diffusely expressed (star) in the mouse lower molar mesenchyme, as in hamster molars. Focused *Dkk1* expression and cusp formation is delayed in the hamster as compared to mouse, for both teeth.

D- Three other WNT and BMP feedback inhibitors show early elevated expression levels in mouse as compared to hamster. Top: transcriptomic profiles in mouse and hamster. Bottom: spatial profile in the bucco-lingual dataset at 15.0 dpc in mouse. The colour scale represents normalised read numbers for each gene. *Sostdc1* is buccal in both mouse molars, while *Grem2* and *Sfrp2* are lingual.

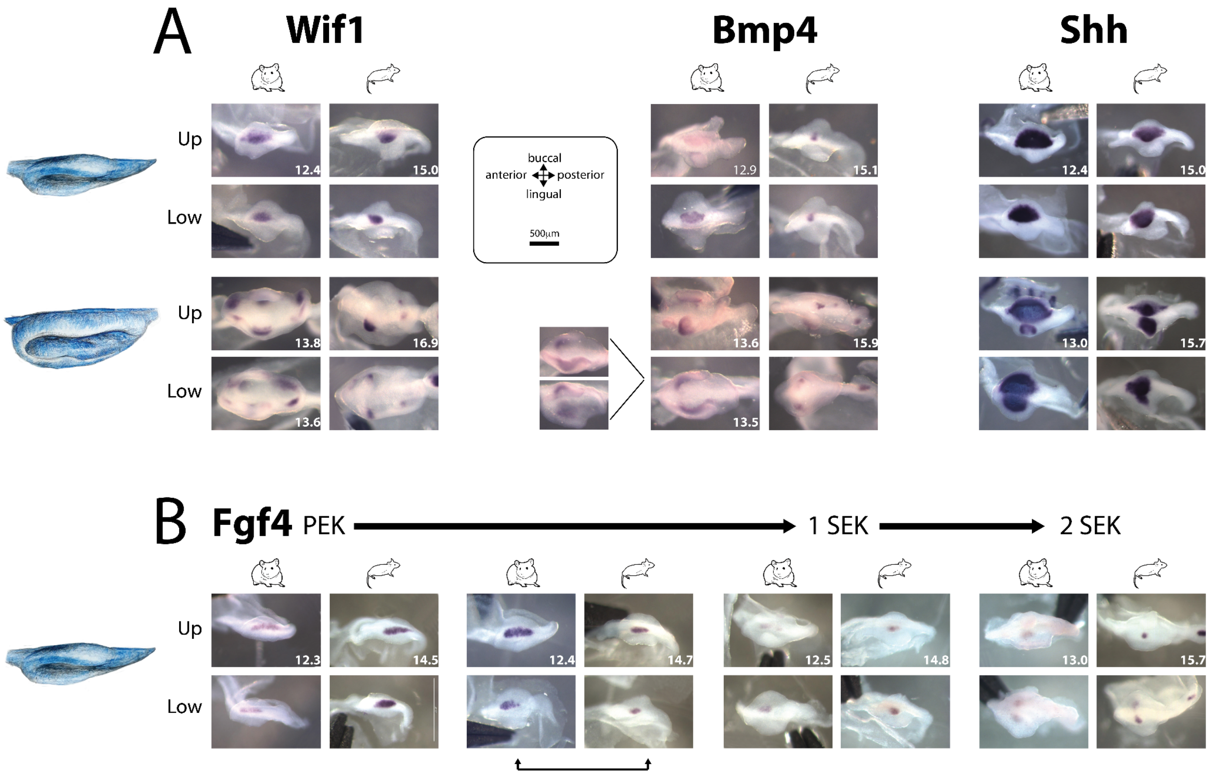


**Figure S10: Mouse molars show a precocious transition to SEK stage, with more focalized expression of signalling molecules.**

A: *Wif1*, *Bmp4* and *Shh* expression territories are more focalized in mouse SEKs. This is seen in early stages with a single SEK (left) or in later stages with more SEKs (right). Mouse and hamster samples are paired for similar advancement of epithelial growth. In hamster, the upper molar is markedly delayed compared to lower molar, therefore when available, we selected slightly older upper samples for the upper molar. Numeric age (days) is indicated in the bottom right of the upper molar picture when lower and upper samples were taken from the same embryo, or in both pictures, when samples were taken from different embryos. For the sake of comparison, *Bmp4* samples shown in Figure 4A are shown next to *Wif1* and *Shh* samples.

B: Transition from the PEK to the 2 SEK stage as seen on tooth germ epithelial parts hybridised against *Fgf4*. Same samples as in Figure 3B, numeric age is added in the corner. Double arrow: The elongated PEK-like expression of *Fgf4* is still visible in hamster, but roundish SEK-like expression of *Fgf4* is already seen in mouse.

Related to Figure 4/

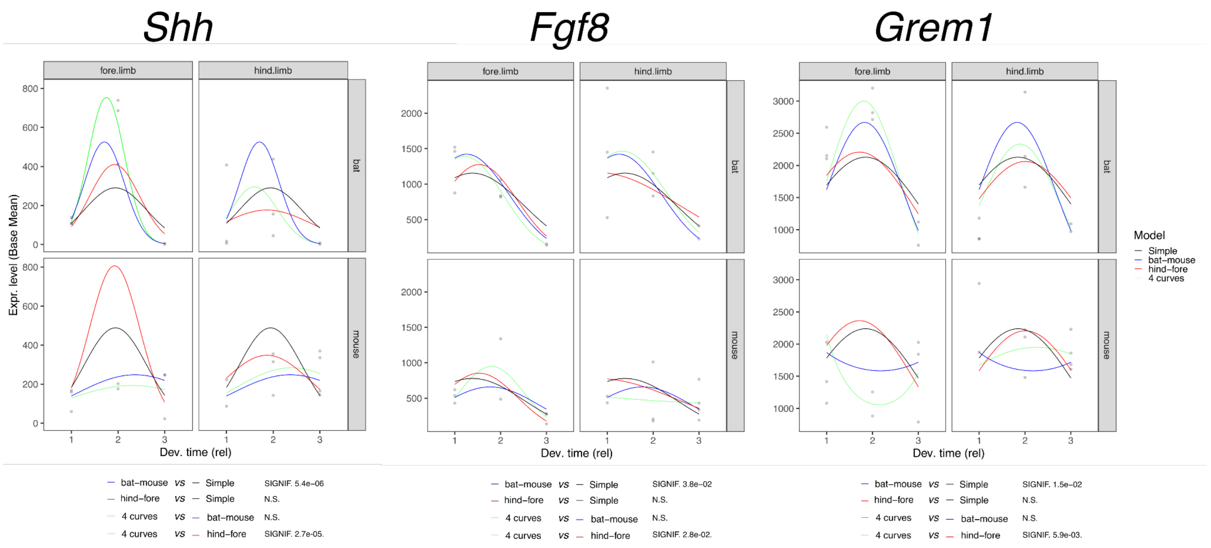


**Figure S11: *Shh*, *Fgf8* and *Grem1* co-evolved in the mouse/bat transcriptome dataset**
For each gene, each quadrant represents expression levels (grey dots) at the 3 timepoints in a given condition (mouse/bat fore/hindlimb), and 4 types of spline modeling: Simple: 1 curve for all conditions; Bat-mouse: 1 curve for the two bat timeseries, 1 curve for the two mouse timeseries; hind-fore: 1 curve for the two hindlimbs, 1 curve for the two forelimbs; 4 curves: 1 curve per condition. Below, we show the results of the statistical tests asking if a first given model fits better than a second model (N.S. non significant).

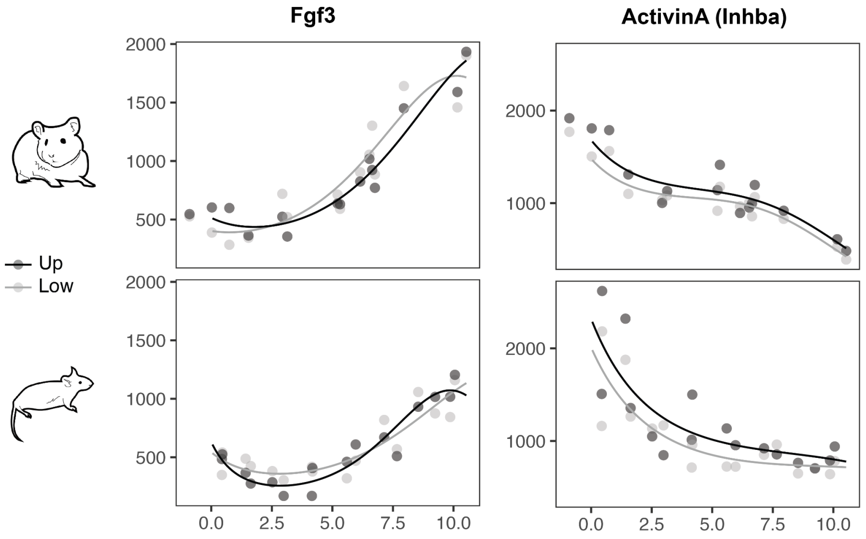


**Figure S12: Transcriptional profiles for *Fgf3* and *Inhba* (coding for ACTIVINβA) suggest that they are not involved in the mouse upper molar evolution.** Below we explain why these genes were potential candidates based on the literature and why we rule them out.
*Fgf3* - It was shown previously that in a heterozygous mutant for the *Fgf3* gene, the first upper molar is smaller and loses its most anterior supplementary cusp, converted into a crest, as seen for the first murine rodents appearing in the paleontological record

(*Potwarmus* genus, (Charles et al., 2009)). In the homozygous mutant, the first upper molar is even smaller and its morphology is severely altered: a longitudinal crest is present, as in murine ancestors or in hamster (the central cusps linked by this crest are malformed), the most anterior cusp is totally suppressed (no crest) but the posterior supplementary cusp is present and large. The overall morphology is thus very far from any ancestral shape, but these shape modifications suggest that making supplementary cusps is developmentally associated with suppressing the longitudinal crest. The molar size is markedly reduced. The suppression of the supplementary cusps is unsurprising, as they form last and therefore will mechanically be the most affected by any reduction of tooth growth. The authors propose that mouse upper molar evolution may have relied on increased FGF3 activity. In our transcriptomes, *Fgf3* showed no increased expression in mouse as compared to hamster (nor in lower/upper mouse molar). The expression levels and profiles are remarkably similar during the cap and bell stage, and hamster expression levels increase more rapidly during the late bell stage than they do in mouse. This is consistent with a role of *Fgf3* in crown growth, the crown being higher in hamster, but suggests that *Fgf3* was not involved in murine dental plan evolution.

*Inhba* - It was shown that adding ActivinβA to the culture medium of lower molars induces the formation of small supplementary cusps on the lingual side (Harjunmaa et al., 2012). In our transcriptomes, the mean expression level and profile are not significantly different between mouse and hamster (nor lower and upper molar in each species).
